# High-throughput organoid screening enables engineering of intestinal epithelial composition

**DOI:** 10.1101/2020.04.27.063727

**Authors:** Benjamin E. Mead, Kazuki Hattori, Lauren Levy, Marko Vukovic, Daphne Sze, Juan D. Matute, Jinzhi Duan, Robert Langer, Richard S. Blumberg, Jose Ordovas-Montanes, Alex K. Shalek, Jeffrey M. Karp

## Abstract

Barrier tissue epithelia play an essential role in maintaining organismal homeostasis, and changes in their cellular composition have been observed in multiple human diseases. Within the small intestinal epithelium, adult stem cells integrate diverse signals to regulate regeneration and differentiation, thereby establishing overall cellularity. Accordingly, directing stem cell differentiation could provide a tractable approach to alter the abundance or quality of specialized cells of the small intestinal epithelium, including the secretory Paneth, goblet, and enteroendocrine populations. Yet, to date, there has been a lack of suitable tools and rigorous approaches to identify biological targets and pharmacological agents that can modify epithelial composition to enable causal testing of disease-associated changes with novel therapeutic candidates. To empower the search for epithelia-modifying agents, we establish a first-of-its-kind high-throughput phenotypic organoid screen. We demonstrate the ability to screen thousands of samples and uncover biological targets and associated small molecule inhibitors which translate to *in vivo*. This approach is enabled by employing a functional, cell-type specific, scalable assay on an organoid model designed to represent the physiological cues of *in vivo* Paneth cell differentiation from adult intestinal stem cells. Further, we miniaturize and adapt the organoid culture system to enable automated plating and screening, thereby providing the ability to test thousands of samples. Strikingly, in our screen we identify inhibitors of the nuclear exporter Xpo1 modulate stem cell fate commitment by inducing a pan-epithelial stress response combined with an interruption of mitogen signaling in cycling intestinal progenitors, thereby significantly increasing the abundance of Paneth cells independent of known WNT and Notch differentiation cues. We extend our observation *in vivo*, demonstrating that oral administration of Xpo1 inhibitor KPT-330 at doses 1,000-fold lower than conventionally used in hematologic malignancies increases Paneth cell abundance. In total, we provide a framework to identify novel biological cues and therapeutic leads to rebalance intestinal stem cell differentiation and modulate epithelial tissue composition via high-throughput phenotypic screening in rationally-designed organoid model of differentiation.

## Introduction

Epithelial barrier tissues enable interaction, exchange, and protection from the external environment. These vital functions are accomplished by specialized epithelial cells integrated with stromal and immune cell populations. Within the small intestine, adult intestinal stem cells (ISCs), conventionally identified by markers including Lgr5 (Barker et al., 2007) and Olfm4 (van der Flier et al., 2009), provide a source of constant regeneration from which an ordered process of differentiation into secretory and absorptive epithelial cells determines composition, thereby steering barrier function. Under homeostatic conditions, WNT, BMP, and Notch signaling actively maintain the ISC niche (Kim et al., 2005; Pinto et al., 2003). However, ISCs have a demonstrated capacity to integrate dietary and immune-derived signals to modulate their self-renewal and differentiation into specific secretory lineages (Beyaz et al., 2016; Biton et al., 2018; von Moltke et al., 2016). Further, following injury, the ISC niche has a remarkable capacity to regenerate from non-stem or quiescent stem pools (Ayyaz et al., 2019; Tetteh et al., 2016; Yan et al., 2017). Cellular identity in the stem cell niche is fluid and responsive to both physiologic and pathologic stimuli (Roulis et al., 2020). In the barrier tissues of the upper respiratory tract and skin, changes in epithelial composition arising from aberrant stem cell differentiation (Naik et al., 2017; Ordovas-Montanes et al., 2018) may be a precipitating factor in infalmmatory diseases. Additionally, shifts in the composition and quality of mature epithelial cells descendant from ISCs are known to occur in the colon and the small intestine of patients suffering from inflammatory bowel disease (Graham and Xavier, 2020; Smillie et al., 2019). Given that barrier tissue stem cells possess significant control over cellular composition and may provide for the restoration of tissue homeostasis in a broad spectrum of human disease, they are a compelling target for therapeutic development.

To support discovery efforts focused on modulating or restoring epithelial barrier composition, there is a need for tools to identify druggable, biological targets involved in regulating epithelial differentiation. Such a tool should employ a physiologically-representative model of the barrier to provide for the best opportunity to identify biological targets that may translate to the *in vivo* context, while also recapitulating cell differentiation processes in a scalable and robustly measurable fashion.

Intestinal organoids—broadly defined as three-dimensional, stem cell-derived, tissue-like cellular structures—have proven to be valuable models of the adult stem cell niche, and preserve known developmental pathways in stem cell differentiation (Sato et al., 2009; Yin et al., 2014). However, because organoid models are dynamic, cellularly and structurally heterogeneous, and typically require complex and costly experimental manipulations, their application in phenotypic high-throughput screens has been limited. Screening activities in organoid models of multiple tissues have so far either focused on clear, genetically-driven behaviors (Dekkers et al., 2013; Korving et al., 2017), or had screening capacity demonstrated on the order of tens of perturbations (Czerniecki et al., 2018). While these are foundational steps towards harnessing organoids for screening, these models have not yet provided for novel biological target discovery which translates to the native *in vivo* tissue.

One approach to adapt organoids for phenotypic high-throughput screening is through the reduction of model complexity around a well-structured hypothesis which incorporates links to *in vivo* tissue biology (Mead and Karp, 2019). Existing chemical tools for small intestinal organoids may enable this approach. The addition of well-characterized small molecules to culture media provides for both enriching intestinal organoids for ISCs, and driving differentiation down specific lineages, by providing physiologically-meaningful cues, such as the modulation of WNT and Notch signaling (Yin et al., 2014). Use of such rationally-directed differentiation has been applied to induce functional Paneth cells from enriched ISCs *in vitro* (Mead et al., 2018). By employing small molecules to mimic known *in vivo* signaling cues of physiological differentiation from a stem cell, it becomes possible to construct organoids representative of lineage-specific stem cell differentiation and frame a screening campaign around identifying new biological targets which may modulate such differentiation. For instance, identification of biological targets and accompanying small molecules which enhance Paneth cell differentiation from ISCs.

Searching for novel targets which enhance Paneth cell differentiation and increase abundance in the native tissue may be therapeutically valuable. Declines in Paneth cell quality and number are observed in inflammatory bowel disease (IBD) (Gassler, 2017; Khor et al., 2011; Liu et al., 2016; McGuckin et al., 2009; Xavier and Podolsky, 2007). Similar Paneth cell aberrations occur in necrotizing enterocolitis (NEC), corresponding with intestinal immaturity and excessive inflammation and systemic infection (McElroy et al., 2013; Sherman et al., 2005; Tanner et al., 2015; White et al., 2017). Emerging evidence suggest that certain viral pathogens, including a subset of coronavirus, may mediate their disruption of the intestinal barrier by targeting and depleting Paneth cells (Wu et al., 2020). Finally, patients with graft versus host disease (GvHD) can exhibit a loss in Paneth cell number and quality, and associated microbial dysbiosis (Eriguchi et al., 2012). Additionally, treatment with R-spondin1, a potentiator of WNT signaling, can elevate the secretion of Paneth-specific alpha-defensins and resolve dysbiosis seen in mice with GvHD by stimulating ISCs to differentiate into Paneth cells (Hayase et al., 2017).

While treatment with R-spondin1 illustrates the importance of stem cell cues driving barrier tissue reconstitution, it faces a major challenge in clinical translation because WNT activation is implicated in precancerous hyperplasia (Han et al., 2017; Okubo and Hogan, 2004; Sansom et al., 2004). Other signaling pathways known to drive Paneth cell differentiation, including Notch signaling, face similar challenges. Activation of Notch signaling amplifies the proliferative progenitor population and promotes an absorptive cell lineage (Fre et al., 2005; Jensen et al., 2000; VanDussen et al., 2012). Conversely, deactivation of Notch signaling amplifies differentiation to all secretory cell types and secretory cell hyperplasia (VanDussen and Samuelson, 2010). A more specific, and niche factor-independent treatment to accomplish ISC-to-Paneth differentiation could provide for safer modulation of tissue composition.

Directing ISCs to preferentially differentiate to Paneth cells independent of known niche signaling (WNT and Notch) offers both a specific hypothesis and a compelling therapeutic target. Accordingly, we have sought to adapt and scale an organoid model to screen for pharmaceutically-actionable biological targets which mediate ISC-to-Paneth differentiation independent of major niche-associated pathways.

## Results

### Small molecule screen for regulators of Paneth cell differentiation

To screen for novel biological targets with established pharmacophores which may regulate Paneth cell differentiation *in vitro* and translate *in vivo*, we require a physiological model system and a scalable screening approach to reasonably scan annotated small molecule space (up to thousands of samples) and measure specific changes of a single cell type (Paneth cells), within a dynamic (differentiation) and heterogeneous (organoid) system. To model physiologically-driven Paneth cell differentiation, we employ a method of small molecule-mediated enrichment and differentiation of intestinal organoids from ISCs to Paneth cells. Murine intestinal organoids are typically expanded in a 3-D Matrigel scaffold with supplemented culture media containing growth factors intended to mimic the ISC niche – epidermal growth factor (E), BMP-antagonist noggin (N), and the aforementioned WNT-pathway enhancer R-spondin1 (R). Spontaneous differentiation of organoids grown in ENR media can be mitigated, and the population of ISCs enriched by the addition of small molecules CHIR99021 (C), an activator of WNT signaling, and valproic acid (V), an activator of Notch signaling. These ISC-enriched cultures can then be differentiated towards Paneth cells with the small molecules (C) and DAPT (D), an inhibitor of Notch signaling, as we have previously shown (Mead et al., 2018; Yin et al., 2014). To measure changes in Paneth cell abundance or quality, we employ a previously demonstrated assay of Paneth cell-specific function and relative-abundance with a commercially-available assay for secreted lysozyme (LYZ) (Mead et al., 2018). To scale this model system and measurement, we adapt conventional 3-D organoid culture into a 2.5-D pseudo-monolayer, where ISC-enriched organoids are partially embedded on the surface of a thick layer of Matrigel (at the Matrigel-media interface), rather than fully encapsulated in the Matrigel structure — an approach similar to others previously reported (Langhans, 2018). This technique enables Matrigel plating, cell seeding, and media additions to be performed in a high-throughput, fully-automated, 384-well plate format and allows for LYZ secretion directly into cell culture media, thereby enhancing measurement sensitivity.

To test our approach, we screened a small molecule library containing well-annotated and specific small molecule inhibitors of a diverse range of biological targets **(Supp. Table 1)** over a 6-day differentiation starting from ISC-enriched organoid precursors (n = 3 biological replicates originating from unique murine donors). Small molecules were added into distinct wells at 4 concentrations per compound (80 nm to 10 μM range) at day 0 and day 3, and at day 6 we measured the functional secretion of LYZ in media supernatants, as a specific marker of Paneth cell enrichment **(Fig. 1A)**. Paneth cell abundance was determined by measuring basally secreted LYZ (LYZ.NS) followed by carbachol (CCh)-induced secretion (LYZ.S), along with cellular ATP as a measure of relative cell number per well. We chose to assay both stimulant-induced (LYZ.S – total cellular LYZ) and basal (LYZ.NS – constitutively secreted LYZ) secretion to distinguish compounds which may mediate changes in Paneth cell quality or secretion (LYZ.NS and LYZ.S uncorrelated) versus changes in Paneth cell abundance (LYZ.NS and LYZ.S correlated). The target-selective inhibitor library contained 433 compounds with high specificity to 184 unique molecular targets—many implicated in stem cell differentiation. In total, our proof-of-concept screen assayed 5,760 unique samples with a 3-measure functional assay.

**Figure 1:**
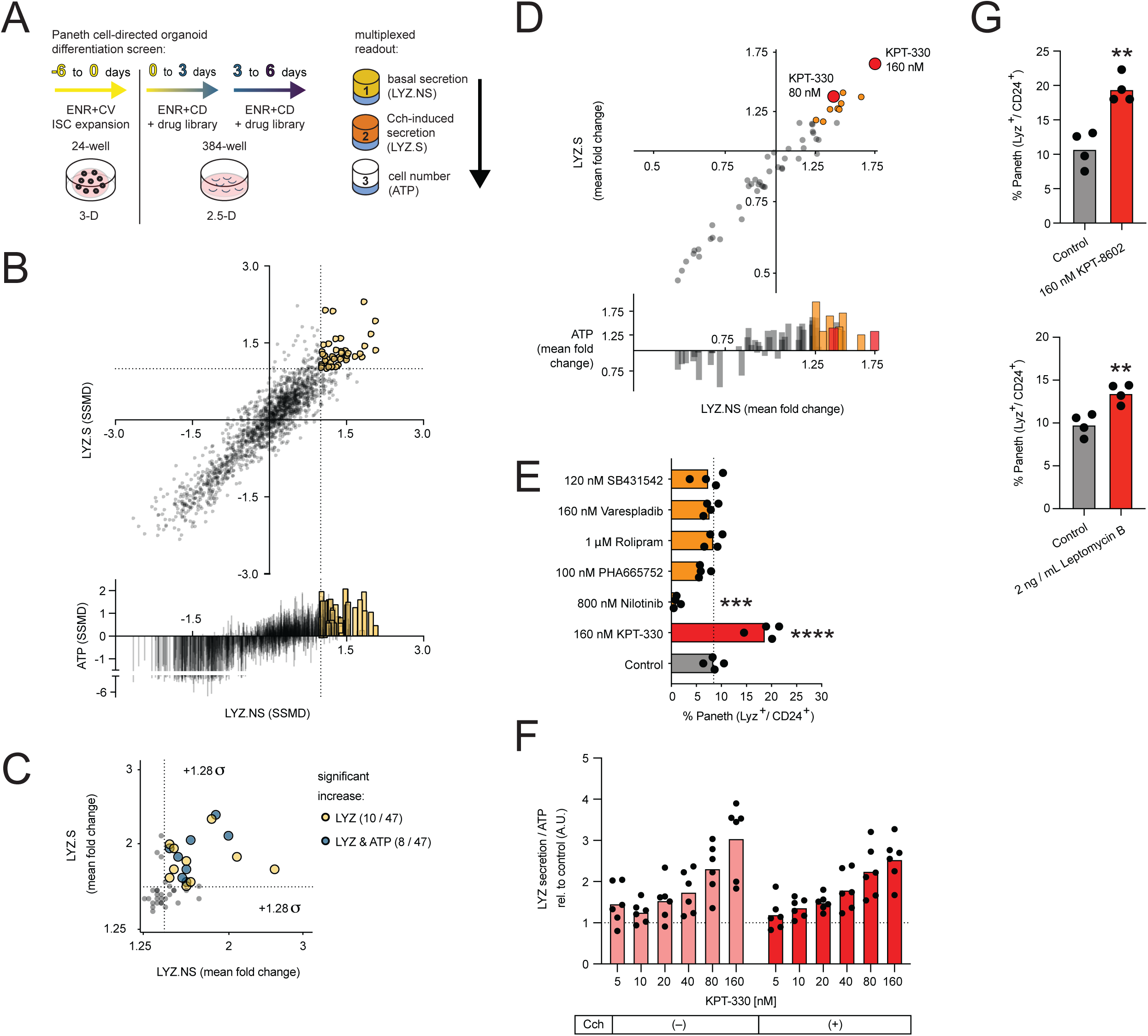
High throughput organoid differentiation screen reveals nuclear export-mediated Paneth cell differentiation. A. Stem-enriched (ENR+CV) to Paneth-enriched (ENR+CD) organoid differentiation screen with a target-selective inhibitor library (433 compounds), assayed with multiplexed functional secretion, both basal and 10uM carbachol-stimulated (Cch) stimulated lysozyme (LYZ) secretion and cell number (ATP) on day 6. B. Replicate strictly standardized mean difference (SSMD) for each assay in primary screen, each point represents the SSMD from 3 replicates of 3 bio. donors relative to whole-plate control, colored points are hits above false positive limit and false negative limit-determined cutoff in both LYZ.NS and LYZ.S assays. C. Mean fold change of assay effect for hits in LYZ.S and LYZ.NS (yellow) or all three assays (blue) in primary screen, only points above 1.28 standard deviations of all treatment mean fold changes for LYZ.S and LYZ.NS are deemed significantly increased. D. Mean fold change for each assay in secondary validation screen (n=8 well replicates, relative to DMSO controls), orange signifies treatments advanced for profiling, red marking most potent compound, KPT-330. E. Flow cytometry for mature Paneth cell fraction of all live cells in 3D-cultured intestinal organoids, treated with 6 hit compounds during 6 days culture in ENR+CD media. Paneth cells identified as lysozyme-positive and CD24-mid cells. Means and individual values are shown (N=4), dotted line represents the average Paneth cell fraction in control samples. One-way ANOVA post-hoc Dunnett’s multiple comparisons test; ****p < 0.0001, ***p < 0.001. F. LYZ secretion assay for organoids differentiated in ENR+CD with increasing concentrations of KPT-330 for 6 days. Organoids were incubated in fresh basal media with or without 10 μM carbachol (Cch) for 3 h on day 6. All data normalized to ATP abundance and standardized to the control in each experiment. Means and individual values are shown (N=6), dotted line represents the control value (1). G. Flow cytometry for mature Paneth cell fraction of all live cells in 3D-cultured intestinal organoids, treated with 160 nM KPT-8602 or 2 ng/mL Leptomycin B during 6 days culture in ENR+CD media. Paneth cells were identified as lysozyme-positive and CD24-mid cells. Means and individual values are shown (N=4). Unpaired two-tailed t-test; **p < 0.01.

We first sought to demonstrate that our screening approach is reproducible, and that our assays measure meaningful function at scale. Following normalization of all measured wells, each assay had an approximate-normal distribution, with lower-value tails corresponding to toxic compounds **(Supp. Fig. 1A)**. Assay values across biological replicates were well correlated, with Pearson correlation values between screen plates ranging from 0.50 to 0.74 **(Supp. Fig. 1B)**. To validate our multiplexed assay’s performance in the screen, we assessed each assay’s performance based on un-treated control wells which are randomly-distributed in each screening plate. Control wells had significantly higher ATP readings than no-cell wells (adj. p < 0.0001), and in the LYZ.NS and subsequent LYZ.S assays, supernatant LYZ was significantly higher in 10 µM CCh-stimulated control wells than in basal control wells (LYZ.NS adj. p < 0.0001, LYZ.S adj. p < 0.05), which in turn were significantly higher than no-cell wells (LYZ.NS adj. p < 0.0001, LYZ.S adj. p < 0.0001) **(Supp. Fig. 1C)**. Following confirmation of reasonable plate reproducibility and assay performance, we next sought to define which molecules meaningfully increased Paneth cell abundance.

We defined primary screen ‘hits’ as having replicate strictly standardized mean differences (SSMDs) in both LYZ.NS and LYZ.S assays greater than the calculated optimal critical value 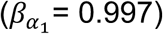 **(Fig. 1B, data in Supp. Table 1, see Methods)**. 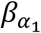 was determined as the intersection minimizing false positive and false negative levels (FPL & FNL error = 0.084) for up-regulation of SSMD-based decisions (Zhang, 2011). The 47 hits correspond to treatment-dose (grouped by biological replicate) combinations that had a statistically significant increase in LYZ.NS and LYZ.S without regard to viability (though most hits per these criteria had positive effects on cellular ATP). Hits were narrowed down to 15 treatment-dose combinations using the z-scored fold-change to select for combinations that elicited a biological effect in the top 10% of values for both LYZ assays relative to the plate (z score > 1.282). Thus, 15 drugs (covering 18 treatment-dose conditions) from 13 unique molecular targets were identified as primary screen hits **(Fig. 1C)**. For molecular targets with more than one hit, only the most robust treatment-dose was selected for further investigation.

To validate primary screen hits against an ENR+CD (not plate) control, while identifying appropriate dose-response ranges and narrowing hits to only the most potent activator(s) of Paneth cell differentiation, we performed a secondary screen with the 13 primary screen hit compounds. Compounds were tested at a narrowed dose range around each treatment’s identified optimal dose from the primary screen (4X below, 2X below and above). Hits in the validation screen were chosen by SSMDs in both LYZ.NS and LYZ.S assays greater than the calculated optimal critical value (β_α1_ = 0.889), with 6 compounds passing this threshold **(Supp. Fig. 1D)**. The same treatment-dose conditions passing the SSMD threshold also had the greatest biological effect, and in particular one compound, KPT-330, a known Xpo1 inhibitor, had two doses representing the greatest and near-greatest biological effect (∼50-75% increases in LYZ.NS and LYZ.S relative to ENR+CD control) **(Fig. 1D)**.

The results of primary and secondary screening reflect a mixture of potential effects which may cause increases in total LYZ secretion. This includes contributions from: (1) enhanced Paneth cell differentiation, (2) altered Paneth cell quality, and (3) changes in total cell number concurrent with differentiation. To better inform how the 6 compounds increased total secreted LYZ, and to isolate only those which enhance Paneth differentiation robustly, we utilized flow cytometry to measure changes in Paneth cell representation within treated organoids during differentiation. Concurrently, to ensure that we do not select for compounds which manifest their behavior only in specific *in vitro* settings, we performed the analyses in the conventional 3-D culture method (4), thus providing control for 2.5-D culture system-specific effects. Single live cells were selected by several gating strategies and Paneth cells were identified as LYZ-high, CD24-mid, side scatter-high (SSC-high) **(Supp. Fig. 1E)**. Only one compound, KPT-330 — the most potent compound in validation screening — significantly enhanced the mature Paneth cell population within differentiating organoids, suggesting KPT-330 induces Paneth differentiation (1) **(Fig. 1E)**. Of the 5 remaining compounds, Nilotinib excluded, none changed organoid composition and are likely driving changes in Paneth quality (2) or are mediating effects dependent on 2.5-D culture (4). Nilotinib significantly decreased Paneth abundance, while significantly increasing total cell number, suggesting the overall increase in bulk LYZ secretion is an effect of increased proliferation (3), or 2.5-D-mediated effect (4).

To examine whether our hits are dependent or independent of canonical stem cell niche signaling, we next evaluated whether the culture media supplements C and D (which mimic physiological Paneth differentiation through WNT activation and Notch inhibition) may alter the effects of our 6 hits. We measured Paneth cell abundance in the canonical ENR culture condition in 3-D (NB because Paneth cells exist in an immature state within ENR, we were unable to robustly quantify Paneth cell number via flow cytometry, and instead used the LYZ secretion assay). This result mirrored our flow cytometry findings in the ENR+CD condition, suggesting that the identified compounds act independently of strong WNT and Notch drivers, and that only KPT-330 is enhancing Paneth cell-specific activity in the conventional organoid culture condition **(Suppl. Fig. 1F)**. Collectively, these results led us to focus solely on KPT-330 and its potential mechanism of Xpo1 inhibition.

### Confirming Xpo1 as a molecular target enhancing Paneth cell differentiation

We next sought to confirm the predicted on-target activity of KPT-330 and investigate the dose-dependency of treatment in enhancing Paneth cell differentiation. KPT-330 is a first-in-class orally-administered FDA-approved drug against multiple myeloma, targeting the nuclear transporter XPO1 (also known as CRM-1).

Administration of KPT-330 below 160 nM for 6 days (NB higher concentrations proved toxic in primary screening) showed LYZ secretion increasing in a dose-dependent manner, with 160 nM of KPT-330 as the most effective dose among tested concentrations **(Fig. 1F)**. To support that Xpo1 is the primary biological target of KPT-330, we used two additional Xpo1 inhibitors: KPT-8602 (a second-generation compound based on KPT-330), and leptomycin B (a canonical Xpo1 inhibitor which acts in a manner distinct from KPT-330 and KPT-8602) (Wang and Liu, 2019). Our flow cytometry results show both KPT-8602 and leptomycin B increasing the proportion of Paneth cells in the organoids **(Fig. 1G)**. LYZ secretion assays with the additional Xpo1 inhibitors show similar Paneth cell-enrichments in both conventional (ENR) and Paneth-differentiating (ENR+CD) culture conditions **(Supp. Fig. 1G, 1H)**. Western blotting for intercellular LYZ per unit weight also confirms enrichment with each of the known XPO1 inhibitors **(Supp. Fig. 1I)**, consistent with the results of LYZ secretion assays and flow cytometry analyses.

KPT-330 is a selective inhibitor of nuclear export (SINE); these molecules act by suppressing the Xpo1-regulated nuclear export of multiple proteins and mRNAs from the nucleus to the cytoplasm — including genes involved in stem cell maintenance and differentiation as well as inflammatory stress response (Sendino et al., 2018). Proteins shuttled by Xpo1 are marked with a nuclear export signal (NES). Additionally, Xpo1 is known to regulate cell cycle through Xpo1’s export-independent role in the regulation of mitosis (Forbes et al., 2015). Based on this evidence, we hypothesized that Xpo1 inhibition might provide for enhanced Paneth cell differentiation by directing ISCs to modulate their differentiation trajectories through alterations in either developmental signaling within the nucleus and / or interfering with cell cycle.

### Longitudinal single-cell RNA-sequencing of differentiation reveals multiple population shifts resulting from Xpo1 inhibition including Paneth cell enrichment

To test the hypothesis that KPT-330 drives Paneth differentiation by altering ISC behavior, we utilized single-cell RNA-sequencing (scRNA-seq) via Seq-Well S^3 (Hughes et al., 2019). We performed a longitudinal comparison between untreated and KPT-330-treated organoids over a 6-day differentiation, with a particular emphasis on early timepoints **(Fig. 2A)**. We collected 17 samples at following timepoints: 6 hours (0.25 days), and 1, 2, 3, 4, or 6 days. Each sample consists of single cells from >1,000 organoids from pre-differentiation ENR+CV organoids and both ENR+CD and ENR+CD + KPT-330 (160 nM) conditions. For time points beyond 2 days, media was refreshed every other day. The resulting dataset consists of 19,877 cells. Unique molecular identifier (UMI), percent mitochondrial, and detected gene distributions are similar across samples, within acceptable quality bounds (genes > 500, UMI < 30,000, percent mitochondrial < 35) **(Supp. Fig. 2A**).

**Figure 2:**
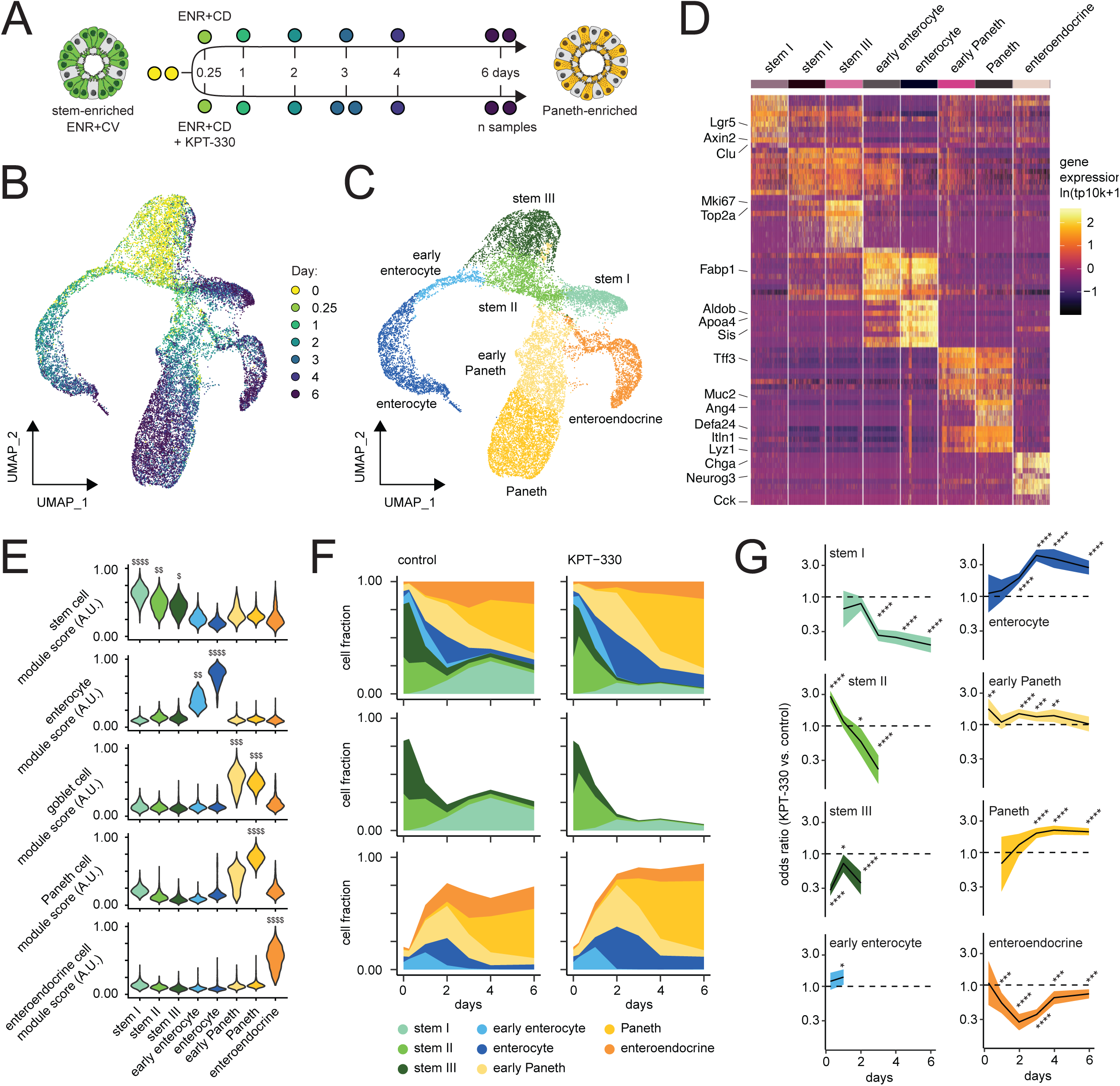
Longitudinal scRNA-seq profiling of Organoid differentiation with Xpo1 inhibition. A. Stem-enriched (ENR+CV) to Paneth-enriched (ENR+CD) organoid differentiation in presence and absence of 160 nM KPT-330, each circle represents a sample of organoids harvested for single-cell RNA-seq over the 6-day time course. B. Organoid differentiation UMAP of all samples labeled by differentiation timepoint. C. Organoid differentiation UMAP of all samples labeled by annotated cell type. D. Log normalized gene expression heatmap for top 10 marker genes by cell type (by log fold change vs. all others). E. Violin plots for all cell types representing module scores derived from gene sets enriched in *in vivo* intestinal stem cells, enterocytes, goblet cells, Paneth cells, and enteroendocrine cells, each score scaled on a range from 0 to 1. Effect size measured as Cohen’s d; $ 0.5 < d < 0.8, $$ 0.8 < d < 1.2, $$$ 1.2 < d < 2, $$$$ d > 2. F. Organoid composition over time between un-treated control and 160 nM KPT-330-treated, for all cell types (top), stem cells (middle), and differentiating cells (bottom). G. Odds ratio enrichment and depletion over differentiation course based on Fisher exact testing with 95% confidence interval for each cell type relative to all others, dotted line at 1. FDR-adjusted Fisher exact testing; *p < 0.05, *p < 0.01, *p < 0.001, *p < 0.0001.

Following normalization, variable feature selection, and principal component dimensional reduction **(see Methods)**, UMAP visualization of the complete dataset reveals the time-course structure along with branches suggestive of distinct lineages arising over the course of differentiation **(Fig. 2B)**. Louvain clustering separated the data into 8 clusters, which we manually annotated **(Fig. 2C)** as mature epithelial and stem cells, based on marker gene expression corresponding to canonical markers of intestinal epithelial cell types **(Fig. 2D & Supp. Table 2)**. Each cluster possessed similar quality metrics, suggesting that clusters are driven by biological and not technical differences **(Supp. Fig. 2B)**. To contextualize and provide a more robust measure of cellular identity of our 8 clusters, we used lineage-defining gene sets from a murine small intestinal scRNA-seq atlas (Haber et al., 2017) to score for enrichment in gene set expression **(Supp. Fig. 2C)**. The eight clusters include three stem-like, two enterocyte, two Paneth, and one enteroendocrine, aligning with our expectation that ENR+CD differentiation should enrich for secretory epithelium cells — principally Paneth and to a lesser extent enteroendocrine **(Fig. 2E)**. To distinguish the three stem-like clusters, and assess physiological relevance, we performed module scoring over gene sets identified to correspond to known ISC subsets *in vivo* (Biton et al., 2018) **(Supp. Fig. 2D)**. We see alignment with the type III and type I ISCs per the nomenclature of Biton et al., along with slight enrichment for a distinct type II **(Supp. Fig. 2E)**, though this population may also be an intermediate between stem I and III populations, sharing markers with both **(Fig. 2D)**. Accordingly, we adopted the naming scheme of Biton et al. to describe the three ISC populations: type I enriched for canonical markers of ISCs (including LGR5), type III distinguished by the high expression of cell cycle genes, and type II appearing as a transitory or intermediate population between I and III.

We next explored changes in cell type representation between organoids treated with KPT-330 versus control. Importantly, in the combined dataset, we do not observe cell clusters unique to KPT-330 treatment, but rather shifts in cluster composition **(Supp. Fig. 2F)**. Both conditions begin with over 75% of cells either stem II or stem III. By day 2, stem I emerges, accounting for approximately 25% of the cells in the control condition, but a smaller proportion in KPT-330-treated organoids. Early enterocytes emerge at day 1, with the continued differentiation to enterocytes, peaking at day 2 and becoming less prevalent by day 4. Early Paneth population appears to crest with enterocytes followed by a transition to Paneth cells continuing to day 6 **(Fig. 2E)**. To better quantify the differences in representation between the KPT-330 and control conditions over time, we performed Fisher exact testing for each cell type relative to all others. This was done for each timepoint where that cell type accounted for at least 1% of cells in both KPT-330 and control samples. We present the relative enrichment or depletion of a cell population with KPT-330 treatment over time as the odds ratio (OR) with a corresponding 95% confidence interval. KPT-330 treatment leads to a depletion of stem I, II, III and enteroendocrine cells over time along with the corresponding enrichment of enterocytes and Paneth cells **(Fig. 2G)**. The observed two-fold enrichment in Paneth cells at day 6 mirrors our flow cytometry observations of a two-fold increase in mature Paneth cells, while also showing the unexpected early enrichment of enterocytes and longer-term depletion of a subset of stem cells — the quiescent stem I population.

### Xpo1 inhibition drives cycling ‘stem II / III’ ISCs into a pro-differentiation state via stress response and suppression of mitogen signaling

Compositional changes during differentiation with KPT-330 are consistent with Xpo1 inhibition acting in a pro-differentiation manner, and our data suggest that the stem II / III populations may be a primary target. In untreated organoids, the expression of *Xpo1* is significantly enriched in the cycling stem III population **(Fig. 3A & Supp. Fig 3A)**, and the expression of genes known to contain a NES (which is required for the nuclear efflux via Xpo1) is enriched in the stem cell populations — most significantly in stem III **(Fig. 3B & Supp. Fig. 3B)** (Fu et al., 2013). More specifically, we know that Xpo1 mediates nuclear signaling processes including the mitogen-activated protein kinase (MAPK) pathway, NFAT, AP-1, and Aurora kinase activity during cell division (Sendino et al., 2018; Sun et al., 2016). With this in mind, we observe the expression of many key mediators in these pathways within the stem populations, and see particular stem III-enrichment in members of MAPK (*Mapk1, Mapk9, Mapk13, Mapk14*), NFAT (*Nfatc3*), AP-1 (*Atf1*), and Aurora kinases (*Aurka, Aurkb*) **(Supp. Fig. 3C)**.

**Figure 3:**
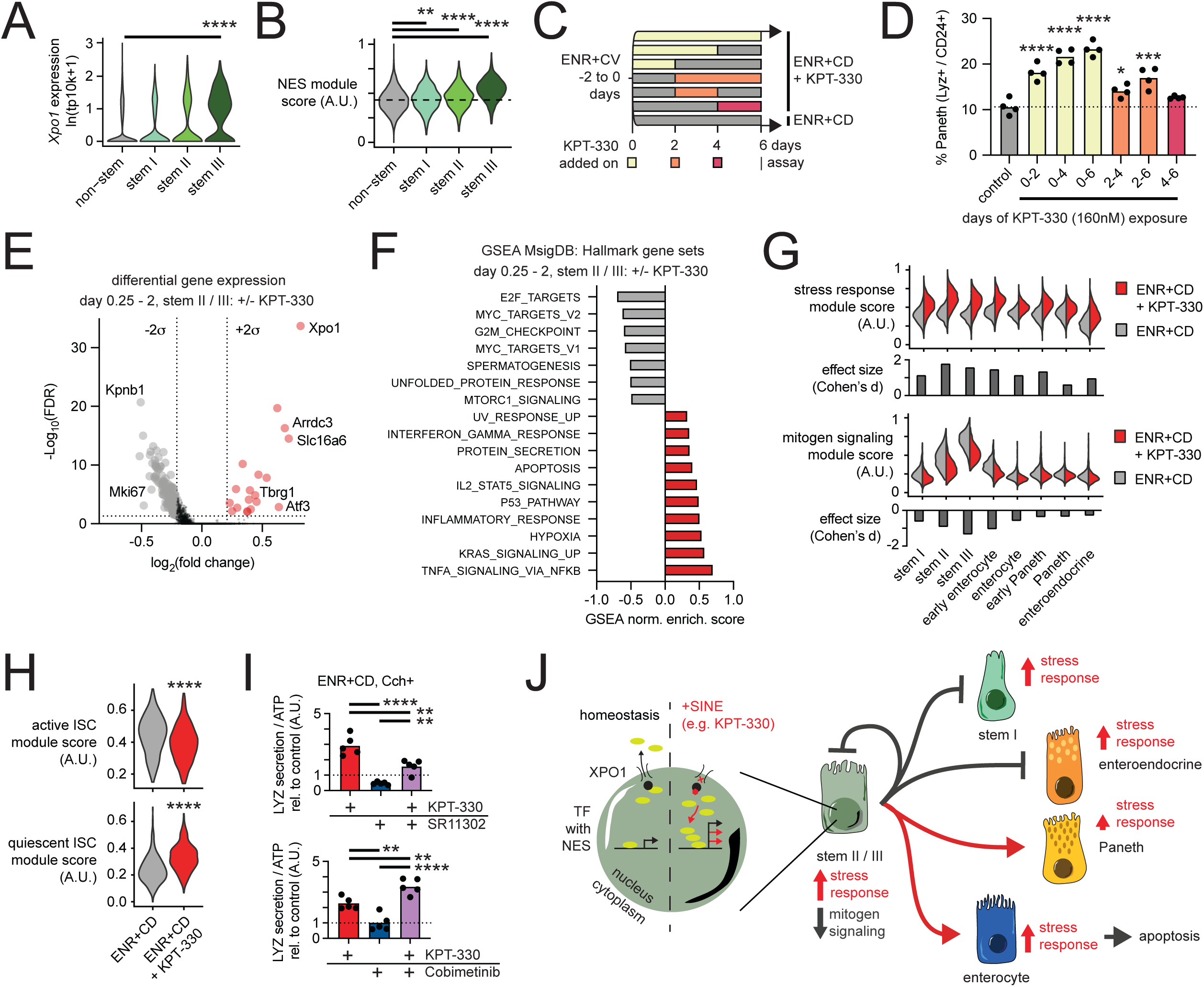
Xpo1 inhibition drives stem cell-specific and pan-epithelial responses to induce differentiation. A. Violin plots of single-cell RNA-seq log normalized (transcripts per 10,000 – tp10k) expression of *Xpo1* in all un-treated control cells split by non-stem, and stem I / II / III annotations. Wilcoxon rank sum test, Bonferroni correction stem I / II / III vs. non-stem; ****p < 0.0001. B. Violin plots of module scores over all cells derived from genes with known nuclear export signals (NES) in all un-treated control cells split by non-stem, and stem I / II / III annotations, each score scaled on a range from 0 to 1. One-way ANOVA post-hoc Dunnett’s multiple comparisons test; ****p < 0.0001, ***p < 0.001, **p < 0.01. C. Time course KPT-330 treatment of ENR+CD differentiating organoids, with treatments over every continuous 2, 4, and 6-day interval. D. Flow cytometry analyses of 3D-cultured intestinal organoids, treated with KPT-330 for the indicated time frame during 6 days culture in ENR+CD media. Paneth cells were identified as lysozyme-positive and CD24-positive cells. Means and individual values are shown (N=4), and the dotted line represents the average of Paneth cell fraction in control samples. One-way ANOVA post-hoc Dunnett’s multiple comparisons test; ****p < 0.0001, ***p < 0.001, *p < 0.05. E. Volcano plot of differentially expressed single-cell RNA-seq log normalized genes between KPT-330-treated and control cells within stem II / III populations in early timepoints (day 0.25-2). Red points are enriched in KPT-330-treated, grey enriched in control. Differential expression based on Wilcoxon rank sum test with significant log2 fold changes based on +/-2 standard deviations of all genes, FDR (Bonferroni correction) cutoff p < 0.05. F. Gene set enrichment analysis (GSEA) normalized enrichment score over all differentially expressed genes between KPT-330-treated and control cells within stem II / III populations in early timepoints (day 0.25-2). Gene sets shown from MSigDB Hallmark v7 with FDR < 0.05, red enriched in KPT-330-treatment, grey enriched in control. G. Split violin plots between KPT-330-treated and control of module scores over all cells derived from significantly enriched (stress response) and depleted (mitogen signaling) genes in KPT-330-treated and control cells within stem II / III populations in early timepoints (day 0.25-2), each score scaled on a range from 0 to 1. Effect size (Cohen’s d) for each module between KPT-330-treated and control within each cell type represented in bar chart below violin plots. H. Violin plots of module scores derived from genes expressed in active and quiescent intestinal stem cells between KPT-330-treated and control cells within stem II / III populations in early timepoints (day 0.25-2), each score scaled on a range from 0 to 1. Two-sided t test; ****p < 0.0001. I. LYZ secretion assay for organoids differentiated in ENR+CD, treated with 10 μM SR11302 (AP-1 inhibitor) or 20 nM Cobimetinib (MEK inhibitor) for 6 days. Organoids were incubated in fresh basal media with or without 10 μM carbachol (Cch) for 3 h on day 6. All data normalized to ATP abundance and standardized to the control in each experiment. Means and individual values are shown (N=5), dotted line represents the control value (1). One-way ANOVA post-hoc Tukey’s multiple comparisons test; ****p < 0.0001, **p < 0.01. J. Proposed mechanism for Xpo1 inhibition driving transcriptional changes manifesting as increased stress responses and reduced mitogen signaling, resulting in rebalanced cycling stem cell fate decisions towards secretory Paneth cells and absorptive enterocytes.

To explore whether the stem II / III population is the principal cellular target of Xpo1 inhibition, we leveraged the dynamic nature of our system and exposed organoids to KPT-330 over every 2, 4, and 6-day interval in the 6-day differentiation and measured final abundance and function of mature Paneth cells at day 6, thereby inferring the relative effect of Xpo1 inhibition on each cell type **(Fig. 3C)**. Of all the 2-day KPT-330 treatments, day 0-2 results in the greatest enrichment in mature Paneth cells, with longer exposure following day 2 providing additional, albeit lesser enrichment. Further, day 2-4 treatment produces moderate enrichment, while day 4-6 is no different than untreated (by flow cytometry) or slightly enriched (by LYZ secretion assay) **(Fig. 3D & Supp. Fig. 3D)**. Using an additional SINE, KPT-8602, we observe similar enrichment behavior as KPT-330 **(Supp. Fig. 3E)**. This data is consistent with Xpo1 inhibition altering stem II / III differentiation – the largest effects of Xpo1 inhibition are concurrent with periods in the differentiation course where stem II / III populations are most abundant. However, this data also suggests that Xpo1 inhibition may not be entirely stem-dependent, given the lesser, but significant increases in Paneth cell number and function with later treatment, where stem II / III populations are greatly diminished.

To better understand the pleiotropic effects of Xpo1 inhibition which may mediate differentiation, we examined the differentially expressed genes between KPT-330 treated and untreated stem II / III populations in the earliest stages of differentiation when they are most abundant (day 0.25-2). Both the most significantly enriched (*Xpo1*) and depleted (*Kpnb1* – a nuclear importin) genes suggest that these cells are significantly impacted by KPT-330 treatment and are enacting changes in expression to reestablish homeostasis of nuclear cargo transit **(Fig. 3E & Supp. Table 3)**. Additional notable genes with significantly increased expression include *Arrdc3* (regulates proliferative processes), *Slc16a6* (principal transporter of ketone bodies; instructional in ISC fate decisions), *Tbgr1* (growth inhibitor), and *Atf3* (regulates stress response in ISCs) (Cheng et al., 2019; Draheim et al., 2010; Jadhav and Zhang, 2017; Zhou et al., 2017). Genes down-regulated by KPT-330 treatment appear related to proliferation and cell cycle, including the marker *Mki67*. In addition to substantial changes within early stem II / III populations, genes regulated by Xpo1 inhibition — including *Xpo1, Atf3, Trp53* (p53), *Ccnd1, Cdk4/6*, and *Cdkn1a* (p21) — have increased expression across all cell types (at all times), but with significant differences in the fraction of cells which express each gene **(Supp. Fig. 3F)**. This suggests that there are both stem II / III -specific responses and pan-epithelial responses to Xpo1 inhibition.

To better contextualize the transcriptional response to KPT-330 treatment in stem II / III cells, we performed gene set enrichment analyses (GSEA) using the v7 molecular signatures database (MSigDB) hallmark collection, which represent specific well-defined biological states or processes across systems (Liberzon et al., 2015; Subramanian et al., 2005). Significant gene sets with FDR < 0.05 reveal two major programs differentially enriched following KPT-330 treatment, with enrichment or depletion quantified through the GSEA normalized enrichment score **(Fig. 3F & Supp. Table 3)**. KPT-330 treatment suppresses programs downstream of mitogen-driven signaling — notably, targets of E2F and MYC, as well as genes involved in cell cycle (G2M checkpoint) — while up-regulating programs broadly resembling a stress response (NFkB signaling, hypoxia, inflammatory response). These responses are in strong agreement with the known effects of Xpo1 inhibition in the context of malignancy.

We next examined whether the responses embodied by the significant differentially-expressed genes in stem II / III (day 0.25-2) may be pan-epithelial or restricted to the cycling stem II / III populations. The stress response module (differentially increased in stem II / III) is substantially increased across all cells during differentiation, with the greatest effect in the stem II / III as well as early mature cell populations, and lowest effect in the mature Paneth cells **(Fig. 3G)**. Conversely, the mitogen signaling module (differentially decreased in stem II / III) is selectively decreased in stem II / III and early enterocyte populations relative to all others. This selectivity corresponds with our observation that the majority of mitogen signaling occurs within the proliferative stem II / III populations relative to the mature populations. As further evidence of altered mitogen signaling impacting stem II / III cells following Xpo1 inhibition, we observe a decrease in a gene module identified by Basak et al. of active ISCs, and a corresponding increase of the quiescent ISC module in our early (day 0.25-2) stem II / III cells **(Fig. 3H)**. Combined with our observation that Xpo1 inhibition blocks the emergence of the quiescent stem I population, our data suggest a model wherein SINE-induced stress response and disruption of mitogen signaling instruct proliferative progenitors to exit cell cycle and differentiate preferentially towards the Paneth and enterocyte lineages, while limiting the accumulation of quiescent stem I cells and enteroendocrine cells.

We sought to clarify this conceptual model with the use of additional small molecule inhibitors known to modulate discrete components of our hypothesized differentiation process, namely: signaling through Xpo1-associated stress response including AP-1 and p53, signaling within the MAPK pathway, and finally Xpo1-mediated effects on mitosis through association with aurora kinases. We began by treating organoids along the ENR+CD differentiation course with SR11302, a small molecule inhibitor of AP-1, to test whether AP-1 is critical to the SINE-induced stress response — both alone and in combination with KPT-330. We observe that SR11302 significantly decreases functional LYZ secretion at the end of the 6-day differentiation, both in combination with KPT-330 and alone **(Fig. 3I)**. This suggests that AP-1 signaling is an important mediator of Paneth differentiation from ISCs.

We next tested whether p53 is a downstream mediator of Xpo1 inhibition by repeating the above assay with two known p53 modulators: a p53 inhibitor pifithrin-α (PFTa), and p53 agonist serdemetan (serd.). Across a wide dose-range, both p53 modulators tested did not alter Paneth cell differentiation — neither alone nor in combination with KPT-330 — suggesting that the KPT-330 stress response is not dependent on p53 signaling modulated by either compound **(Supp. Fig. 3G)**. With the same assay, we began to probe the mitogen signaling response by adding the MEK inhibitor, cobimetinib (shown by Basak et al. to induce the quiescent ISC population), in combination with KPT-330. Interestingly, cobimetinib alone did not significantly alter Paneth cell differentiation, but gave a different result in combination with KPT-330 **(Fig. 3I)**. We next sought to test whether the regulation of cell cycle via mitogen signaling may be an important downstream mediator following Xpo1 inhibition. Inhibition of Cdk4/6 with palbociclib both alone and in combination with KPT-330 did not alter Paneth cell differentiation **(Supp. Fig. 3H)**, but inhibition of aurora kinase b with ZM447439 did significantly increase Paneth cell differentiation (notably, ZM447439 was also a lower-effect size hit in our primary screen) **(Supp. Fig. 3I)**. Combined, these experiments suggest that the SINE-induced stress response may be mediated by AP-1 but not p53, while suppression of mitogen signaling is not dependent on ERK, but is further enhanced by ERK inhibition. Additionally, the non-exporter-related action of Xpo1 during cell cycle (which interacts with aurora kinase) may further contribute to the observed pro-differentiation effect.

In total, our analyses suggest that Xpo1 inhibition drives Paneth cell enrichment through the modulation of cell state within cycling ISCs (stem II / III). Further, this modulation includes a confluence of pan-epithelial stress response and suppression of mitogen signaling within stem II / III. We observe the cycling stem population becoming transiently quiescent, thereby favoring differentiation towards the Paneth and enterocyte lineages (the latter being a short-lived population relative to the former) over a more balanced transition to the mature lineages and the quiescent stem pool (stem I) **(Fig. 3J)**.

### Low dose oral Xpo1 administration *in vivo* induces selective expansion of the Paneth cell compartment

Based on our understanding of Xpo1 inhibition in stem-enriched organoids, we hypothesized that SINE compounds may selectively enrich the epithelium for Paneth cells *in vivo*. Our findings in organoids suggest that SINE treatment is independent of the niche cues of WNT and Notch, and acts specifically on cycling stem cells (which are abundant in the epithelial crypts). While Xpo1 inhibition may enrich both for Paneth cells and enterocytes, by virtue of the relatively long Paneth cell lifespan (Ireland et al., 2005) we would expect a longer-term accumulation of Paneth cells *in vivo* relative to enterocytes. Additionally, because Xpo1 inhibition in organoids does not expand the stem cell pool but rather rebalances patterns of differentiation, we expect an increase in Paneth cell number following SINE treatment *in vivo* to be restricted to the spatial constraints of non-hypertrophic crypts and proportional to the initial number of cycling progenitors. This suggests *that in* vivo increases in Paneth cell number may be modest, thus requiring a particularly sensitive method of quantification.

Following a similar protocol as previously reported for SINE treatment in the context of cancer (Arango et al., 2017; Azmi et al., 2013; Hing et al., 2016; Zheng et al., 2014), KPT-330 was administered at a dose of 10 mg/kg via oral gavage every other day over a two-week span in C57BL/6 wild-type mice, and body weight was monitored for any clear toxicity. Within the treatment group, we observed significant weight loss indicative of toxicity **(Supp. Fig. 4A)**. Given animal weight loss on the standard chemotherapeutic dosage regimen, and additional evidence that sustained dosage of SINEs adversely impacts T cell populations (Tyler et al., 2017), we sought to explore dosing regimens well below 10 mg/kg, to assess if a pro-Paneth phenotype may exist below potential toxicities.

We repeated the two-week study with oral gavage of KPT-330 every other day at doses corresponding to 50-fold (0.2 mg/kg), 200-fold (0.05 mg/kg), and 1,000-fold (0.01 mg/kg) decrease in the 10 mg/kg dose conventionally used in a cancer setting. Because Paneth cell number and quality is known to physiologically change along the length of the small intestine, and diseases associated with Paneth cells most frequently present distally (Abraham and Cho, 2009), we sought to profile how Xpo1 inhibition may differentially affect proximal and distal small intestine. We tracked animal weight every other day and collected the proximal and distal thirds of the small intestine at day 14 for histological quantification of Paneth, stem, and goblet populations **(Fig. 4A)**. In this lower-dose regimen, we observe no significant changes in animal weight, suggesting that we are outside the gross toxicity range **(Supp. Fig. 4B)**. Paneth cells were counted within well preserved crypts — with at least 30 crypts quantified per animal (representative images **Supp. Fig. 4C) —** and the counts were averaged. Compellingly, within this lower dose regimen, we observe significant increases in Paneth cell abundance both in the proximal and distal small intestine at doses of 0.01 mg/kg, and proximally at 0.2 mg/kg. We additionally quantified the abundance of Olfm4+ stem cells as well as PAS+ goblet cells within the same animals to ascertain whether the effect of SINE treatment was restricted to the Paneth cell compartment (representative images **Supp. Fig. 4D**,**E)**. Interestingly, we observe a significant increase in Olfm4+ stem cells within the distal SI at doses of 0.01mg/kg corresponding to the group with the greatest increase in Paneth cells **(Fig. 4C)**, suggesting a potential expansion of the stem cell niche commensurate with increased Paneth cell abundance. We did not identify any significant changes in the developmentally-related goblet cell population **(Fig. 4D)**. In total, this data suggests that SINE-treatment may be a meaningful approach to specifically increase Paneth cell abundance *in vivo*, and further validates our framework for using models of organoid differentiation in small molecule screening.

**Figure 4:**
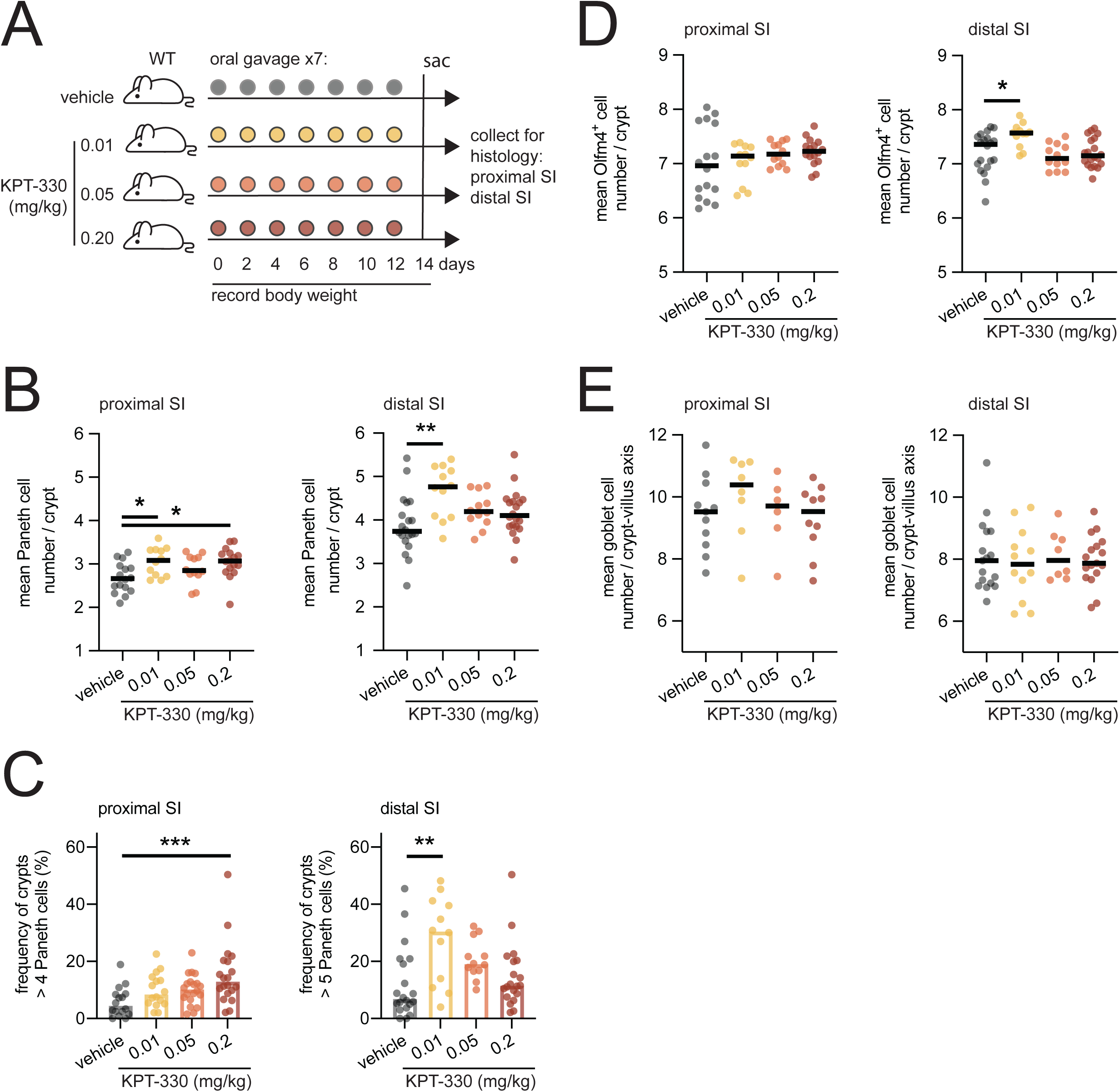
Xpo1 inhibition with KPT-330 increases Paneth cell number *in vivo*. A. Design for *in vivo* oral gavage of KPT-330 in wild-type (WT) C57BL/6 mice. B. Mean Paneth cell number per crypt in proximal or distal third of small intestine, quantified by blinded histological counting. Means and individual values are shown. N=12 (0.01 and 0.05 mg/kg, proximal), N=20 (vehicle and 0.2 mg/kg, distal), N=12 (0.01 and 0.05 mg/kg, distal). One-way ANOVA post-hoc Dunnett’s multiple comparison test; **p < 0.01, *p < 0.05. C. Frequency of crypts with 4 or more (proximal third) or 5 or more (distal third) Paneth cells per animal across KPT-330 treatment groups, from blinded histological counts. Means and individual values are shown. N=16 (vehicle and 0.2 mg/kg, proximal), N=12 (0.01 and 0.05 mg/kg, proximal), N=20 (vehicle and 0.2 mg/kg, distal), N=12 (0.01 and 0.05 mg/kg, distal). One-way ANOVA post-hoc Dunnett’s multiple comparisons test; ***p < 0.001, **p < 0.01. D. Mean Olfm4^+^ stem cell number per crypt in proximal or distal third of small intestine, quantified by blinded histological counting. Means and individual values are shown. N=16 (vehicle and 0.2 mg, proximal), N=12 (0.01 and 0.05 mg/kg, proximal), N=20 (vehicle and 0.2 mg/kg, distal), N=12 (0.01 and 0.05 mg/kg, distal). One-way ANOVA post-hoc Dunnett’s multiple comparison test; *p < 0.05. E. Mean PAS^+^ goblet cell number per crypt-villus axis in proximal or distal third of small intestine, quantified by blinded histological counting. Means and individual values are shown. N=11, 8, 6, 10 (vehicle, 0.01, 0.05. 0.2 mg/kg, proximal), N=17, 12, 8, 17 (vehicle, 0.01, 0.05. 0.2 mg/kg, distal).

## Discussion

Here, we demonstrate that, by employing phenotypic small molecule screening in a rationally and physiologically motivated organoid model, we can uncover novel biological targets and clinically-relevant small molecules which translate to and inform *in vivo* tissue stem cell biology. Further, this approach to small molecule phenotypic screening enables a specific, functional readout in a dynamic and heterogeneous organoid model. Our approach provides perturbation capacity nearly two orders of magnitude greater than existing examples of non-genetic perturbations in organoid models, thereby enabling screens within the space of annotated small molecule libraries and empowering novel biological target discovery.

By using a model that focuses on differentiation to a specific lineage (the Paneth cell), we are able to resolve a pathway and compounds which direct ISC fate decisions to drive subtle but significant effects on the *in vivo* tissue. We identify a series of compounds — known to inhibit the nuclear exporter Xpo1 — acting on cycling ISCs by inducing a program of stress response and decreased mitogen signaling. This ISC response re-balances self-renewal and differentiation towards Paneth cell differentiation. Recent work on mitogen and stress response control of re-entry into cell cycle may provide important context on the necessity of overlap of these two responses (Yang et al., 2017). Specifically, mother cells will transmit p53 protein and *Ccnd1* transcripts to daughter cells, which, based on the abundance of transmitted signal, will either immediately re-enter cell cycle or commit to a quiescent state. Transitions between quiescence and proliferation within the ISC niche have important roles in tissue homeostasis and regeneration. Quiescent pools of crypt-residing or adjacent cells serve as reserve populations which upon injury-dependent depletion of cycling stem cells will re-establish cycling progenitors and maintain homeostatic tissue regeneration (Ayyaz et al., 2019; Yousefi et al., 2017). Further, a transient quiescent ISC state facilitates secretory enteroendocrine cell differentiation (Basak et al., 2017), and may explain why we see even synergistic Paneth cell enrichment with a combination of ERK and Xpo1 inhibition. Further, ERK inhibition may affect differentiation by either suppression of enterocyte differentiation (De Jong et al., 2016) or by augmenting Wnt signaling (Heuberger et al., 2014) — both alternate explanations for the observed synergy. Additionally, we see that the pan-epithelial stress response induced by Xpo1 inhibition *in vitro* involves the AP-1 pathway — which also appears to play an important role in Paneth differentiation — and may be mediated via transcriptional changes in Atf3. *In vivo*, Atf3 is implicated in the regulation of stress responses in disease of the barrier tissue (Glal et al., 2018; Zhou et al., 2017), where Xpo1 may be one way to access these observed responses for therapeutic use.

Because Xpo1 inhibition is inherently pleiotropic, there are many possible leads to consider as downstream mediators within ISCs driving the observed shift in fate decision making. Future work exploring these many complex interacting effects is warranted. Additionally, exploring the potential local cellular niches of what we describe as stem II/III within these perturbed and evolving organoids may provide great clarity on the extent to which the organoid stem niche does and does not recapitulate *in vivo*. As previously shown, events of symmetry-breaking are well-modeled within intestinal organoids and spatial structuring is important in cell fate decisions (Serra et al., 2019). While tools to profile these kinds of changes are presently limited, they will become increasingly available. Studies of this nature will better illuminate biological targets which may better translate to large tissue effects *in vivo*.

While we have demonstrated that Xpo1 inhibition *in vivo* significantly increases the abundance of Paneth cells within the small intestinal crypts, we believe there are two key mediators of biological potency which may explain heterogeneity of response along the small intestine, and may be investigated to enhance effect in future studies. The first is a consideration of the pharmacological profile and tissue distribution of the agent we used — KPT-330, and the second are elements of tissue and stem cell niche biology which our organoid model does not presently incorporate. Future studies may improve the potency of Xpo1 inhibition within the small intestinal crypts by re-formulating or targeting the delivery of SINE compounds to the crypt microanatomy and favor distal over proximal small intestine, where disease biology is most relevant and we observe the most robust expansion of Paneth cells. Additionally, as our knowledge of the *in vivo* stem cell niche improves, organoid models can be refined in an iterative fashion, further enhancing model fidelity and increasing the probability of compound translation from *in vitro* to *in vivo*.

This approach – employing a physiologically-motivated set of cues to modulate intestinal stem cell differentiation and layering on a screen for new regulators of that differentiation at scale – may be further applied both within the small intestine and more broadly across adult barrier tissues to modulate the composition of epithelial specialists through novel molecular targets and associated small molecules. For example, this same framework may be applied in the context of enteroendocrine cell development within the small intestine to explore ways in which important hormone secretion — including gpl-1 — may be modulated. While establishing the appropriate model and a screening assay for a study of enteroendocrine differentiation is not trivial, it should be possible based on the approach we have demonstrated here. Overall, we provide a framework to construct organoid models of lineage-specific differentiation that can uncover pathways regulating differentiation and reveal compounds controlling barrier tissue composition.

## Acknowledgements

This work was supported in part by the Koch Institute Support (core) Grant P30-CA14051 from the National Cancer Institute. We thank the Koch Institute Swanson Biotechnology Center for technical support, specifically Jaime Cheah and Christian Soule with the High Throughput Science facility, the Flow Cytometry facility, and the Histology facility. In addition, we thank the Broad Institute Walk-Up Sequencing core for providing Illumina NovaSeq services, and the Harvard Digestive Disease Center and NIH grant P30DK034854, for the provision of reagents.

B.E.M. was supported by the National Science Foundation graduate research fellowship program and the Massachusetts Institute of Technology – GlaxoSmithKline (MIT-GSK) Gertrude B. Elion Postdoctoral fellowship. K.H. received fellowships from The Japanese Biochemical Society (The Osamu Hayaishi Memorial Scholarship for Study Abroad), Mochida Memorial Foundation for Medical and Pharmaceutical Research, and The Uehara Memorial Foundation. J.O.M was supported by the Richard and Susan Smith Family Foundation, the HHMI Damon Runyon Cancer Research Foundation Fellowship (DRG-2274-16), the AGA Research Foundation’s AGA-Takeda Pharmaceuticals Research Scholar Award in IBD – AGA2020-13-01, and the HDDC Pilot and Feasibility P30 DK034854. R.L was supported by the NIH (DE013023). R.S.B. was supported by the NIH (DK088199). A.K.S. was supported by the Beckman Young Investigator Program, the Pew-Stewart Scholars Program for Cancer Research, a Sloan Fellowship in Chemistry, the NIH (1DP2GM119419, 1U54CA217377). J.M.K. was supported by the Kenneth Rainin Foundation Innovator and Breakthrough awards, the Crohn’s and Colitis Foundation (#624458), and the NIH (HL095722).

## Author Contributions

Conceptualization, B.E.M., K.H., A.K.S., J.M.K.;

Methodology, B.E.M., L.L., D.S., K.H., J.D.M., J.D., R.S.B.;

Software, B.E.M., D.S.;

Formal Analysis, B.E.M., K.H., D.S.;

Investigation, K.H., B.E.M., L.L., M.V.;

Resources, J.M.K., A.K.S., R.L., R.S.B.;

Writing – Original Draft, B.E.M., K.H., L.L., D.S., R.L., J.O.M., A.K.S., J.M.K.;

Writing – Review & Editing, B.E.M., K.H., M.V., J.D.M., R.L., J.O.M., A.K.S., J.M.K.;

Visualization, B.E.M., K.H., D.S., J.O.M., A.K.S., J.M.K.;

Supervision, J.M.K., R.L., J.O.M., A.K.S.;

Project Administration, B.E.M.;

Funding Acquisition, B.E.M., K.H., J.O.M., R.S.B., R.L., A.K.S., J.M.K.

## Methods

### Experimental Model: Mice

All studies were performed under animal protocols approved by the Massachusetts Institute of Technology (MIT) Committee on Animal Care. Proximal and / or distal small intestine was isolated from wild-type C57BL/6 mice of both sexes, aged between one and six months in all experiments.

### Crypt isolation and culture

Small intestinal crypts were isolated as previously described^23^. Briefly, the small intestine was harvested, opened longitudinally, and washed with ice-cold Dulbecco’s Phosphate Buffer Saline without calcium chloride and magnesium chloride (PBS0) (Sigma-Aldrich) to clear the luminal contents. The tissue was cut into 2-4 mm pieces with scissors and washed repeatedly by gently pipetting the fragments using a 10-ml pipette until the supernatant was clear. Fragments were rocked on ice with crypt isolation buffer (2 mM EDTA in PBS0; Life Technologies) for 30 min. After isolation buffer was removed, fragments were washed with cold PBS0 by pipetting up and down to release the crypts. Crypt-containing fractions were combined, passed through a 70-μm cell strainer (BD Bioscience), and centrifuged at 300 rcf for 5 min. The cell pellet was resuspended in basal culture medium (2 mM GlutaMAX (Thermo Fisher Scientifc) and 10 mM HEPES (Life Technologies) in Advanced DMEM/F12 (Invitrogen)) and centrifuged at 200 rcf for 2 min to remove single cells. Crypts were then cultured in a Matrigel culture system (described below) in small intestinal crypt medium (100X N2 supplement (Life Technologies), 100X B27 supplement (Life Technologies), 1 mM N-acetyl-L-cysteine (Sigma-Aldrich) in basal culture medium) supplemented with differentiation factors at 37°C with 5% CO_2_. Pen/strep (100X) was added for the first four days of culture post-isolation only.

Small intestinal crypts were cultured as previously described^23^. Briefly, crypts were resuspended in basal culture medium at a 1:1 ratio with Corning™ Matrigel™ Membrane Matrix – GFR (Fisher Scientific) and plated at the center of each well of 24-well plates. Following Matrigel polymerization, 500 μl crypt culture medium (ENR+CV) containing growth factors EGF (50 ng/ml, Life Technologies), Noggin (100 ng/ml, PeproTech) and R-spondin 1 (500 ng/ml, PeproTech) and small molecules CHIR99021 (3 μM, LC Laboratories or Selleckchem) and valproic acid (1 mM, Sigma-Aldrich) was added to each well. ROCK inhibitor Y-27632 (10 μM, R&D Systems) was added for the first two days of ISC culture only. Cell culture medium was changed every other day. After 4 days of culture, crypt organoids were expanded as and enriched for ISCs under the ENR+CV condition. Expanding ISCs were passaged every 4-6 days in the ENR+CV condition.

### Organoid culture, differentiation, and passaging

After 2 to 6 days of culture under ENR+CV condition, ISCs were differentiated to Paneth cells. Briefly, ISC culture gel and medium were homogenized via mechanical disruption and centrifuged at 300 rcf for 3 min at 4°C. Supernatant was removed and the pellet resuspended in basal culture medium repeatedly until the cloudy Matrigel was almost gone. On the last repeat, pellet was resuspended in basal culture medium, the number of organoids counted, and centrifuged at 100 rcf for 1 min at 4°C. The cell pellet was resuspended in basal culture medium at a 1:1 ratio with Matrigel and plated at the center of each well of 24-well plates (∼100-250 organoids/well). Following Matrigel polymerization, 500 μl crypt culture medium (ENR+CV) was added to each well. Cell culture medium was changed every 2-4 days depending on seeding density.

### High-throughput screening

For 384-well plate high-throughput screening, ISC-enriched organoids were passaged and split to single cells with TyrpLE (Thermo Fisher Scientific) and cultured for 2-3 days in ENR+CVY prior to transfer to a “2.5D” 384-well plate culture system. To prepare for “2.5D” plating, cell-laden Matrigel and media were homogenized via mechanical disruption and centrifuged at 300 rcf for 3 min at 4°C. Supernatant was removed and the pellet washed and spun in basal culture medium repeatedly until the cloudy Matrigel above the cell pellet was gone. On the final wash, pellet was resuspended in basal culture medium, the number of organoids counted, and the cell pellet was resuspended in ENR+CD medium at ∼7 clusters/μL. 384-well plates were first filled with 10 μL of 70% Matrigel (30% basal media) coating in each well using a Tecan Evo 150 Liquid Handling Deck, and allowed to gel at 37°C for 5 minutes. Then 30 μL of cell-laden media was plated at the center of each well of 384-well plates with the liquid handler, and the plates were spun down at 100 rcf for 2 minutes to embed organoids on the Matrigel surface. Compound libraries were pinned into prepped cell plates using 50 nL pins into 30 μL media/well. Cells were cultured at 37°C with 5% CO_2_ for six days in ENR+CD medium supplemented with the tested compounds with a media change at three days. On day six, lysozyme secretion and cell viability were assessed using Lysozyme Assay Kit (EnzChek) and CellTiter-Glo 3D (CTG 3D) Cell Viability Assay (Promega), respectively, according to the manufacturers’ protocols. Briefly, screen plates were washed 3x with FluoroBrite basal media (2 mM GlutaMAX and 10 mM HEPES in FluoroBrite DMEM (Thermo Fisher Scientific)) using a BioTek 406 plate washer with 10 min incubations followed by a 1 min centrifugation at 200 rcf to settle media between washes. After removal of the third wash, 30 μL of non-stimulated FluoroBrite basal media was added to each screen well using a Tecan Evo 150 Liquid Handling Deck from a non-stimulated treatment master plate, and plates were incubated for 30 min at 37°C. After 30 minutes, the top 15 μL of media from each well of the screen plate was transferred to a non-stimulated LYZ assay plate containing 15 μL of 20X DQ LYZ assay working solution using a Tecan Evo 150 Liquid Handling Deck. The non-stimulated LYZ assay plate was covered, shaken for 10 min, incubated for 50 min at 37°C, then fluorescence measured (shake 10 s; 494 mm/518 nm) using a Tecan M1000 Plate Reader. After the media transfer to the non-stimulated LYZ assay plate, the remaining media was removed from the screen plate and 30 μL of Stimulated FluoroBrite basal media (supplemented with 10 μM CCh) was added to each screen well using a Tecan Evo 150 Liquid Handling Deck from a stimulated treatment master plate, and plates were incubated for 30 min at 37°C. After 30 minutes, the top 15 μL of media from each well of the screen plate was transferred to a stimulated LYZ assay plate containing 15 μL of 20X DQ LYZ assay working solution using a Tecan Evo 150 Liquid Handling Deck. The stimulated LYZ assay plate was covered, shaken for 10 min, incubated for 50 min at 37°C, then fluorescence measured (shake 10 s; 494 mm/518 nm) using a Tecan M1000 Plate Reader. Finally, 8 μL of CTG 3D was added to each well of the screen plate, which was shaken for 30 min at room temperature, then luminescence read (shake 10 s; integration time 0.5-1 s) to measure ATP.

Primary screens were performed using the Target Selective Inhibitor Library (Selleck Chem). Assays were performed in triplicate using four compound concentrations (0.08, 0.4, 2, and 10 μM).

### Screen Analysis

A custom R script and pipeline was used for analysis of all screen results. Results (excel or .csv files) were converted into a data frame containing raw assay measurements corresponding to metadata for plate position, treatments, doses, cell type, and stimulation. Raw values were log_10_ transformed, then a LOESS normalization was applied to each plate and assay to remove systematic error and column/row/edge effects using the formula (Mpindi et al., 2015):

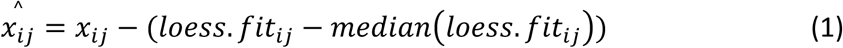

where 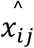 is the loess fit result, *x*_*ij*_ is the log_10_ transformed value at row *i* and column *j*, and *loess. fit*_*ij*_ is the value from loess smoothed data at row *i* and column *j* calculated using R loess function with span 1.

Following LOESS normalization, a plate-wise fold change (FC) calculation was performed on each well to normalize plates across the experiment. This was calculated by subtracting the median of the plate (as control) from the LOESS normalized values:

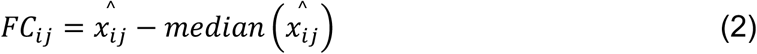

Replicate strictly standardized mean difference (SSMD) was used to determine the statistical effect size of each treatment in each assay (treatment and dose grouped by replicate, n=3) relative to the plate using the formula for the robust uniformly minimal variance unbiased estimate (UMVUE) (Zhang, 2011):

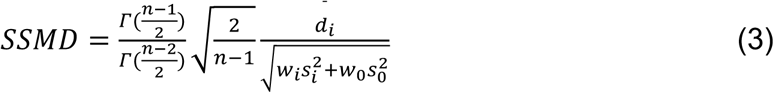

where 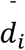 and *s*_*i*_ are respectively the sample mean and standard deviation of *d*_*ij*_s where *d*_*ij*_ is the FC for the *i*th treatment on the *j*th plate. *Γ*(·) is a gamma function. 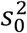 is an adjustment factor equal to the median of all 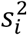 s to provide a more stable estimate of variance. *w*_*i*_ and *w*_*0*_ are weights equal to 0.5 with the constraint of *w*_*i*_ + *w*_*0*_ = 1. *n* is the replicate number.

Mean FC (the arithmetic mean of all samples grouped by treatment and dose across replicates) was used to determine the z-score for each treatment and dose with the formula:

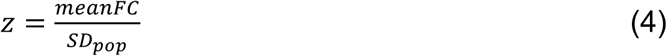

where *SD*_*pop*_ is the standard deviation of all mean-FC’s.

All calculated statistics were combined in one finalized data table and exported as a .csv file for hit identification. A primary screen “hit” was defined as having SSMDs for both LYZ assays greater than the optimal critical value 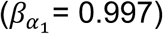 and being in the top 10% of a normal distribution of FC values for both assays with a z-score cutoff > 1.282. 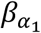 was determined by minimizing the false positive (FPL) and false negative (FNL) levels for up-regulation SSMD-based decisions by solving for the intersection of the formulas (Zhang, 2011):

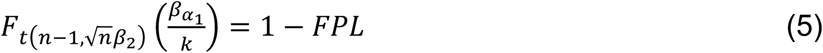

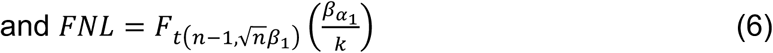

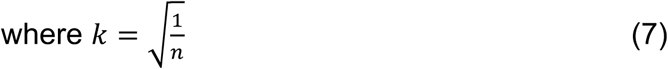

where 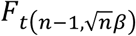 (·) is the cumulative distribution function of non-central *t*-distribution 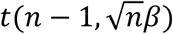 and *n* is the number of replicates, *β*_2_ is a SSMD bound for FPL of 0.25 (at least very weak effect), and *β*_1_ is a SSMD bound for FNL of 3 (at least strong effect).

Hit treatments were thus selected to have a well-powered statistical effect size as well as a strong biological effect size. Optimal dose per hit treatment was determined by SSMD for both LYZ assays.

### Secondary lysozyme secretion assay screen

Confirmatory secondary screening with primary hits was performed using the above 384-well plate method. The screen was conducted with 4-plate replicates with a base media of ENR+CD. Media was supplemented with compound at day 0 and day 3 (n=8 well replicates per dose) at four different doses: two-fold above, two-fold below, and four-fold below the optimal final dose for each respective treatment. Additionally, each plate carried a large number of ENR+DMSO or ENR+CD+DMSO (vehicle) control wells (n=100 for ATP, and n=25 for LYZ.NS and LYZ.S) for robust normalization. ATP, non-stimulated lysozyme activity and CCh-stimulated lysozyme activity was again measured and the collected data was again processed in a custom R-script, per primary screen with slight modification. Values were log_10_ transformed, and a plate-wise FC was calculated for each well based on the median value of ENR+CD+DMSO (vehicle) control wells to normalize plate to plate variability. The following formula was used:

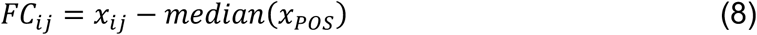

Where *x*_*ij*_ is the log_10_ transformed value at row *i* and column *j*, and *x*_*POS*_ are the values of the positive control wells. For the ATP assay, all vehicle-only wells were used as the control. For the LYZ.NS assay, non-stimulated vehicle only wells were used. For the LYZ.S assay, vehicle only wells that were non-stimulated in the LYZ.NS assay then stimulated in the LYZ.S were used.

Once normalized, the replicate SSMD was calculated using formula (3) to quantify statistical effect size with 8 replicate differences taken relative to the respective plate ENR+DMSO or ENR+CD+DMSO median value. A primary hit was considered validated when SSMDs for both LYZ assays was greater than the optimal critical value 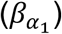 of 0.889. 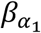 was determined using formula (5) with an FPL error of 0.05 for a more stringent cut off, FNL was not considered. Optimal doses were chosen for treatments with multiple validated doses by taking the most potent (highest mean fold change relative to ENR+CD control) dose in both LYZ assays.

### Lysozyme secretion assay

ISC-enriched organoids in 3-D Matrigel culture were passaged to a 48- or 96-well plate and cultured with ENR or ENR+CD media containing DMSO or each drug for 6 days. DMSO- or drug-containing media were changed every other day. On day 6, cells were washed with basal media twice and treated with basal media with or without 10 μM carbachol for 3 hours in a CO2 incubator at 37°C. Conditioned media was collected and used for lysozyme assay (Thermo, E-22013) following the manufacturer’s instruction. The fluorescence was measured using excitation/emission of 485/530 nm. CTG 3D Reagent was added afterward, and the cell culture plate was incubated on an orbital shaker at RT for 30 min to induce cell lysis and to stabilize the luminescent signal. The solution was replaced to a 96-well white microplate, and luminescent signals were measured by a microplate reader (infinite M200, Tecan). The standard curve was prepared by diluting recombinant ATP (Promega, P1132). For both assays, a polynomial cubic curve was fitted to a set of standard data, and each sample value was calculated on the Microsoft Excel.

### Flow cytometry

ISC-enriched organoids in 3-D Matrigel culture were passaged to a 24- or 48-well plate and induced differentiation for 6 days by ENR+CD media containing DMSO or each drug indicated in the figures. DMSO- or drug-containing media were changed every other day. On day 6, cells were washed twice with basal media, then harvested from Matrigel by the mechanical disruption in TrypLE Express (Thermo, #12605010) to remove Matrigel and dissociate organoids to single cells. After vigorous pipetting and incubation at 37°C for 15 min, the cell solution was diluted twice with basal media and centrifuged at 300 rcf for 3 min. The cell pellet was resuspended in FACS buffer (PBS containing 2% FBS) and replaced into a 96-well Clear Round Bottom Ultra-Low Attachment Microplate (Corning, #7007). The cell solution was centrifuged again at 300 rcf for 3 min at 4°C to pellet the cells. Cells were stained with Zombie-violet dye (BioLegend, # 423113) at 100X for viability staining for 20 min at RT in the dark. After centrifugation for 3 min at 300 rcf, cells were fixed in fixation buffer (FACS buffer containing 1% formaldehyde (Thermo, #28906)) for 15 min on ice in the dark. Cells were centrifuged again for 3 min at 300 rcf and blocked with staining buffer (FACS buffer containing 0.5% Tween20 (Sigma, P2287)) for 15 min at RT in the dark. Pelleted cells by the centrifugation for 3 min at 300 rcf are stained with FITC-conjugated anti-lysozyme antibody (Dako, F0372) and APC-conjugated anti-CD24 antibody (Biolegend, #138505) at 100X for 45 min at RT in the dark. The cell pellet was washed once with FACS buffer, resuspended in FACS buffer, and filtered through 5 mL test tube with cell strainer snap cap (Corning, #352235). Flow cytometry was performed using an LSR Fortessa (BD; Koch Institute Flow Cytometry Core at MIT). Flow cytometry data were analyzed using FlowJo X v10.6.1 software.

### Western blotting

Organoid-containing gel was homogenized in basal medium and centrifuged at 300 rcf for 3 min. Organoid pellet was lysed with ice-cold Pierce IP Lysis Buffer (Thermo Fisher Scientific, #87787) containing Halt Protease Inhibitor Cocktail, EDTA-Free (Thermo Fisher Scientific, #87785) and incubated on ice for 20 min. The lysate was centrifuged at 17,000 rcf for 10 min, and the supernatant was combined with NuPAGE LDS Sample Buffer (Thermo Fisher Scientific, NP0007). Protein concentration was determined by Pierce 660 nm Protein Assay (Thermo Fisher Scientific, #22660) and normalized to the lowest concentration among each sample set. Samples were incubated at 70°C for 10 minutes and resolved by SDS-PAGE using NuPAGE 4-12% Bis-Tris Protein Gels (Thermo Fisher Scientific) followed by electroblotting onto Immun-Blot PVDF Membrane (Biorad, 1620174) using Criterion Blotter with Plate Electrodes (Biorad, #1704070). The membranes were blocked with 2% Blotting-Grade Blocker (Biorad,1706404) in TBS-T (25 mM Tris–HCl, 140 mM NaCl, 3 mM Potassium Chloride and 0.1% Tween 20) and then probed with appropriate antibodies, diluted in TBS-T containing 2% BSA (Sigma, A7906) and 0.05% sodium azide (Sigma, #71289). The primary antibody against lysozyme was purchased from Abcam (ab108508). HRP-linked anti-rabbit IgG antibodies were purchased from Cell Signaling Technology (#7074). Chemiluminescent signals were detected by LAS4000 (GE Healthcare) using Amersham ECL Select Western Blotting Detection Reagent (GE Healthcare, #45-000-999), and total protein signals were obtained by Odyssey Imaging System (LI-COR Biosciences) using REVERT Total Protein Stain Kit (LI-COR Biosciences, #926-11010).

### Animal study

8-10 weeks old wild type C57BL/6NCrl male mice (#027) were purchased from Charles River. Mice were housed under 12 h light/dark cycle and provided food and water ad libitum. 0.01, 0.05, 0.2 or 10 mg/kg of KPT-330 were injected orally using a disposable gavage needle (Cadence Science, #9921) at 10 μL/g weight. KPT-330 was dissolved in DMSO initially and further diluted in sterile PBS containing Pluronic F-68 Non-ionic Surfactant (Gibco, #24040032) and Polyvinylpyrrolidone (PVP, Alfa Aesar, A14315, average M.W. 58,000); the final concentration of DMSO is 2%, Pluronic is 0.5%, and PVP is 0.5%. KPT-330 was administered every other day for two weeks, 7 injections in total (days 0, 2, 4, 6, 8, 10, 12), and mice were sacrificed at day 14. All animal studies are approved by the Committee on Animal Care (CAC) at Massachusetts Institute of Technology.

### Histology

The small intestine (SI) was collected from mice and divided into three parts. Only proximal and distal SI were kept in PBS, and medial SI was discarded. Each SI was opened longitudinally and washed in PBS. SI was rolled using the Swiss-rolling technique and incubated in 10% Neutral Buffered Formalin (VWR, 10790-714) for 24 h at RT. Fixed tissues were embedded in paraffin, and 4 μm sections were mounted on slides. For immunohistochemistry, slides were deparaffinized, antigen retrieved using heat-induced epitope retrieval at 97°C for 20 min using citrate buffer pH 6, and probed with appropriate antibodies followed by DAB staining. An antibody against lysozyme was purchased from Abcam (ab108508), Ki67 from BD Biosciences (#550609), and Olfm4 from Cell Signaling Technology (#39141). For McManus Periodic Acid Schiff (PAS) reaction, slides were deparaffinized, oxidized in periodic acid, and stained with Schiff reagent (Poly Scientific, s272) followed by counterstaining with Harris Hematoxylin. Slides were scanned by Aperio Slide Scanner (Leica) and cells were counted on Aperio eSlide Manager. Slides were blinded and randomized before counting, and all cell types were counted in all well-preserved crypts along the longitudinal crypt-villus axis.

### Single-cell RNA-sequencing and alignment

A single-cell suspension was obtained from organoids cultured under either ENR+CD or ENR+CD + 160 nM KPT-330 for the differentiation time course as detailed in **Fig. 2A**. Briefly, organoids at each sampling were harvested from 4-6 pooled Matrigel domes, totaling >1,000 organoids per sample. Excess Matrigel was removed per previously described washing protocol, and organoids were resuspended in TrypLE at 37C for 15 min, with vigorous homogenization through a p200 pipette tip every 5 min. After 15 min, the suspension was passed through a 30 uM cell strainer twice, and counted under brightfield microscopy with trypan blue staining for viable single cells.

We utilized Seq-Well S^3 for massively parallel scRNA-seq, for which full methods are published (Hughes et al., 2019) and available on the Shalek Lab website (www.shaleklab.com). Briefly, ∼15-20,000 cells were loaded onto a functionalized-polydimethylsiloxane (PDMS) array preloaded with ∼80,000 uniquely-barcoded mRNA capture beads (Chemgenes; MACOSKO-2011-10). After cells had settled into wells, the array was then sealed with a hydroxylated polycarbonate membrane with pore sizes of 10 nm, facilitating buffer exchange while confining biological molecules within each well. Following membrane-sealing, buffer exchange across the membrane permits cell lysis, mRNA transcript hybridization to beads, and bead removal before proceeding with reverse transcription. The obtained bead-bound cDNA product then underwent Exonuclease I treatment (New England Biolabs; M0293M) to remove excess primer before proceeding with second strand synthesis.

Following Exonuclease I treatment, the beads mixed with 0.1M NaOH for 5 min at room temperature to denature the mRNA-cDNA hybrid product on the bead. Second strand synthesis was performed with a mastermix consisting of 40uL 5x maxima RT buffer, 80uL 30% PEG8000 solution, 20uL 10mM dNTPs, 2uL 1mM dn-SMART oligo, 5uL Klenow Exo-, and 53ul of DI ultrapure water, which was added to the beads and incubated for 1 hour at 37°C with end-over-end rotation. After second strand synthesis, PCR amplification was performed using KAPA HiFi PCR Mix (Kapa Biosystems KK2602). Specifically, a 40uL PCR Mastermix consisting of 25 uL of KAPA 5X Mastermix, 0.4 uL of 100 uM ISPCR oligo, and 14.6 uL of nuclease-free water was combined with 2,000 beads per reaction. Following PCR amplification, whole transcriptome products were isolated through two rounds of SPRI purification using Ampure Spri beads (Beckman Coulter, Inc.) at both 0.6X and 0.8x volumetric ratio and quantified using a Qubit.

Sequencing libraries were constructed from whole transcriptome product using the Nextera Tagmentation method on a total of 800 pg of pooled cDNA library per sample. Tagmented and amplified sequences were purified through two rounds of SPRI purification (0.6x and 0.8x volumetric ratios) yielding library sizes with an average distribution of 500-750 base pairs in length as determined using the Agilent hsD1000 Screen Tape System (Agilent Genomics). Arrays were sequenced within multi-sample pools on an Illumina Nova-Seq through the Broad Institute walk-up sequencing core. The read structure was paired end with Read 1 starting from a custom read 1 primer containing 20 bases with a 12bp cell barcode and 8bp unique molecular identifier (UMI) and Read 2 being 50 bases containing transcript information. Sequencing read alignment was performed using version 2.1.0 of the Drop-seq pipeline previously described in (Macosko et al., 2015). For each Nova-Seq sequencing run, raw sequencing reads were converted from bcl files to FASTQs using bcl2fastq based on Nextera N700 indices that corresponded to individual samples. Demultiplexed FASTQs were then aligned to the mm10 genome using STAR and the DropSeq Pipeline on a cloud-computing platform maintained by the Broad Institute. Individual reads were tagged with a 12-bp barcode and 8-bp unique molecular identifier (UMI) contained in Read 1 of each sequencing fragment. Following alignment, reads were grouped by the 12-bp cell barcodes and subsequently collapsed by the 8-bp UMI for digital gene expression (DGE) matrix extraction and generation.

### Single-cell RNA-sequencing analysis

Prior to analysis, DGE matrices were pre-processed to remove cellular barcodes with less than 500 unique genes, greater than 35% of unique molecular identifiers (UMIs) corresponding to mitochondrial genes, low outliers in standardized house-keeping gene expression (Tirosh et al., 2016), barcodes with greater than 30,000 UMIs, and cellular doublets identified through manual inspection and use of the DoubletFinder algorithm (McGinnis et al., 2019). These pre-processed DGEs are deposited as GEO GSE148524.

After quality and doublet correction, we performed integrated analysis on a combined dataset of 19,877 cells, with quality metrics for gene number, captured UMIs, and percent mitochondrial genes reported in **Fig. S2**. To better control for potential batch effects that may arise in sample handling and library preparation, dimensional reduction and clustering was performed following normalization with regularized negative bionomical regression as implemented in Seurat V3 via SCTransform (Hafemeister and Satija, 2019). We performed variable gene identification and dimensionality reduction utilizing the first 9 principal components based on the elbow method to identify 8 cell type clusters using Louvain clustering (Resolution = 0.45). Following UMAP visualization, we used log-normalized RNA expression for all differential gene expression tests, gene set enrichment analyses, and gene module scoring. We identified genes enriched across clusters using the Wilcoxon rank sum test, with genes expressed in at least 20% of cells, and a minimum log-fold change of 0.5, to identify generic cell types, and corroborated these cell type identities relative to gene signatures coming from an established murine small intestinal scRNA-seq atlas (Haber et al., 2017). Gene modules were scored within each cell based on enrichment in gene set expression relative to randomly selected genes of comparable expression levels in each cell (Tirosh et al., 2016), via the AddModuleScore function within Seurat v3. In addition to cell-type module scoring from Haber et al., we incorporated gene sets for ISC sub-typing from (Biton et al., 2018), in addition to gene sets representing ISC activity (Basak et al., 2017), and genes known to contain NES from the ValidNESs database (Fu et al., 2013).

To quantify enrichments in cell populations between treatment and control within the dataset, we utilized Fisher’s exact test for each cell type relative to all others at each timepoint. We only considered populations for testing where that cell type accounted for at least 1% of cells in both KPT-330 and control samples. We present the relative enrichment or depletion of a cell population with KPT-330 treatment over time as the odds ratio (OR) with a corresponding 95% confidence interval, and FDR-adjusted p values with significance as ‘*’’s denoted in corresponding figure legend.

Gene set enrichment analysis (GSEA) was performed on the full rank-ordered list of differentially-expressed genes (without fold-change or p value cutoffs) using the piano R package (Väremo et al., 2013), and the MsigDB hallmark v7 gene sets (Liberzon et al., 2015; Subramanian et al., 2005). Gene sets with at least 25 and no more than 500 matching genes were considered, and only gene sets with an FDR-corrected p value of < 0.05 were retained.

### Quantification and statistical analysis

Methodology for statistical analysis of screens is detailed above. For each subsequent experiment, replicate type and number are reported in corresponding figure legends, along with statistical tests performed and either ‘$’ classifications for Cohen’s D effect sizes, or ‘*’ classifications for p values.

## Data Availability Statement

Supplemental information described below are available upon request. The single-cell RNA sequencing data reported in this paper will be made available shortly through GEO.

## Supplemental Information

**Supplementary Table 1**

Results from primary and secondary lysozyme (LYZ) secretion screening grouped by compound and dose, reported as log2 fold change (FC), standard z score, and strictly standardized mean difference (SSMD).

**Supplementary Table 2**

Marker genes from organoid differentiation time course single-cell RNA-seq, as determined by Wilcoxon differential expression testing of cluster versus rest.

**Supplementary Table 3**

Differentially expressed genes and gene set enrichment analysis (GSEA) over differentially expressed genes between KPT-330 treated and untreated stem II / III cells over days 0.25, 1, 2 in organoid differentiation time course single-cell RNA-seq.

## Figures & Legends

**Supplementary Figure 1.**
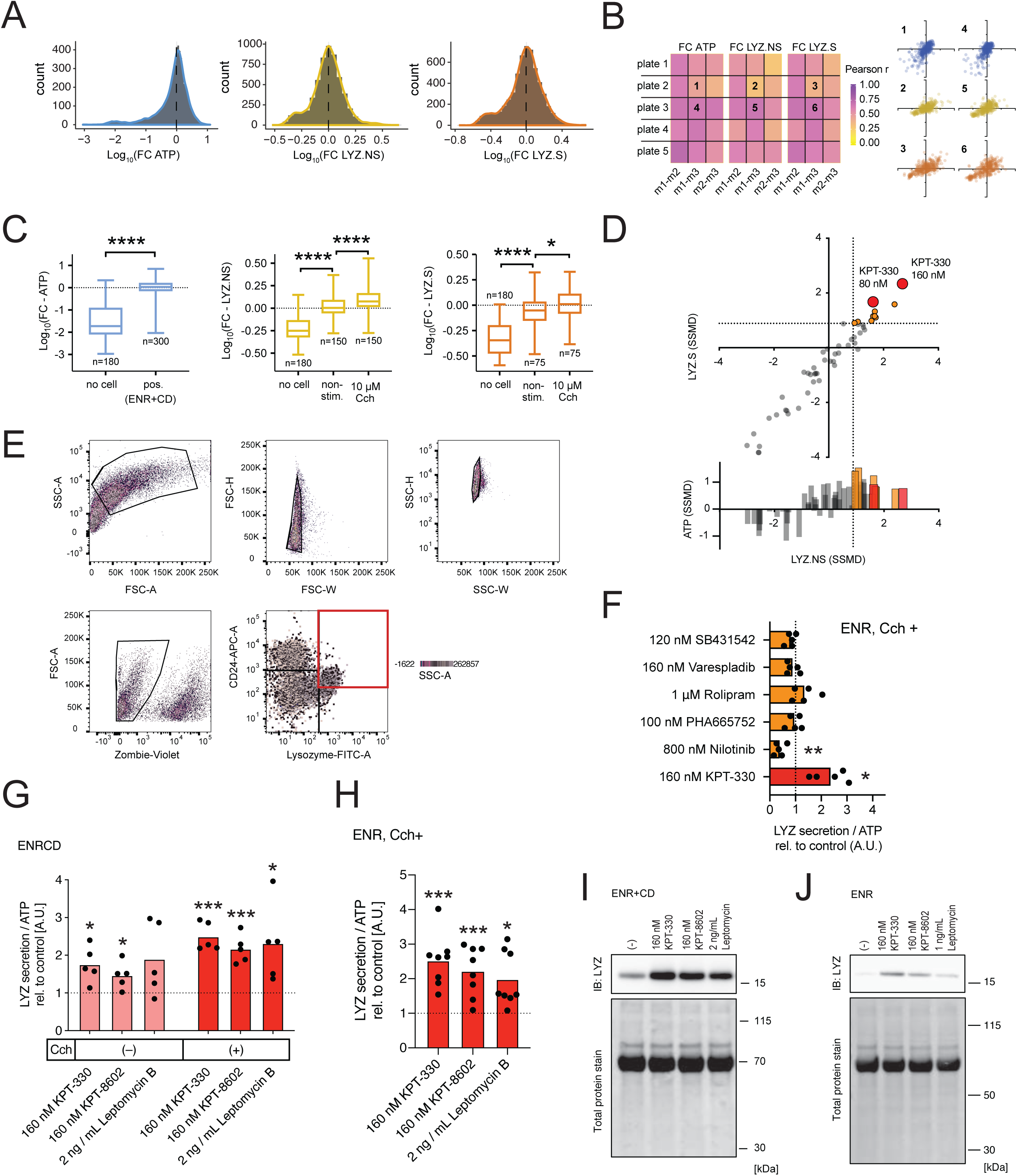
A. Distributions of all sample data (n=5676 wells) for each assay following data transformation and normalization, dotted line indicates median of distribution from which fold change calculations are determined. B. Pearson correlation (r) between all sample wells by screen plate and biological replicate (m1, m2, m3), with representative correlation plots shown for assay plates 1 through 6. C. ATP, LYZ.NS, LYZ.S assay controls across all plates and replicates, Welch’s t test for ATP, one-way ANOVA post-hoc Dunnett’s multiple comparison test * adj. p<0.05, **** adj. p< 0.0001. D. Replicate strictly standardized mean difference (SSMD) for each assay in secondary validation screen, each point represents the SSMD from 8 well-replicates relative to DMSO control, orange signifies treatments passing cutoffs in both LYZ.NS and LYZ.S assays, red marking most potent compound, KPT-330. E. Flow cytometry gating strategy to select viable mature Paneth cells, final gate outlined in red. F. LYZ secretion assay for organoids differentiated in ENR with 6 hit compounds for 6 days. Organoids were incubated in fresh basal media with 10 μM carbachol (Cch) for 3 h on day 6. All data normalized to ATP abundance and standardized to the control in each experiment. Means and individual values are shown (N=5), dotted line represents the control value (1). One sample t-test compared to 1, followed by the two-stage linear step-up method of Benjamini, Krieger and Yekutieli for adjusting p-values; **p < 0.01, *p < 0.05. G. LYZ secretion assay for organoids differentiated in ENR+CD with 160 nM KPT-330, 160 nM KPT-8602 or 2 ng/mL Leptomycin B for 6 days. Organoids were incubated in fresh basal media with or without 10 μM carbachol (Cch) for 3 h on day 6. All data normalized to ATP abundance and standardized to the control in each experiment. Means and individual values are shown (N=5), dotted line represents the control value (1). One sample t-test compared to 1, followed by the two-stage linear step-up method of Benjamini, Krieger and Yekutieli for adjusting p-values; ***p < 0.001, **p < 0.01, *p < 0.05. H. LYZ secretion assay for organoids differentiated in ENR with 160 nM KPT-330, 160 nM KPT-8602 or 2 ng/mL Leptomycin B for 6 days. Organoids were incubated in fresh basal media with 10 μM carbachol (Cch) for 3 h on day 6. All data normalized to ATP abundance and standardized to the control in each experiment. Means and individual values are shown (N=5), dotted line represents the control value (1). One sample t-test compared to 1, followed by the Two-stage linear step-up method of Benjamini, Krieger and Yekutieli for adjusting p-values; ***p < 0.001, *p < 0.05. I. Western blotting of intracellular LYZ in 3D-cultured intestinal organoids, cultured in ENR+CD media for 6 days. J. Western blotting of intracellular LYZ in 3D-cultured intestinal organoids, cultured in ENR media for 6 days.

**Supplementary Figure 2.**
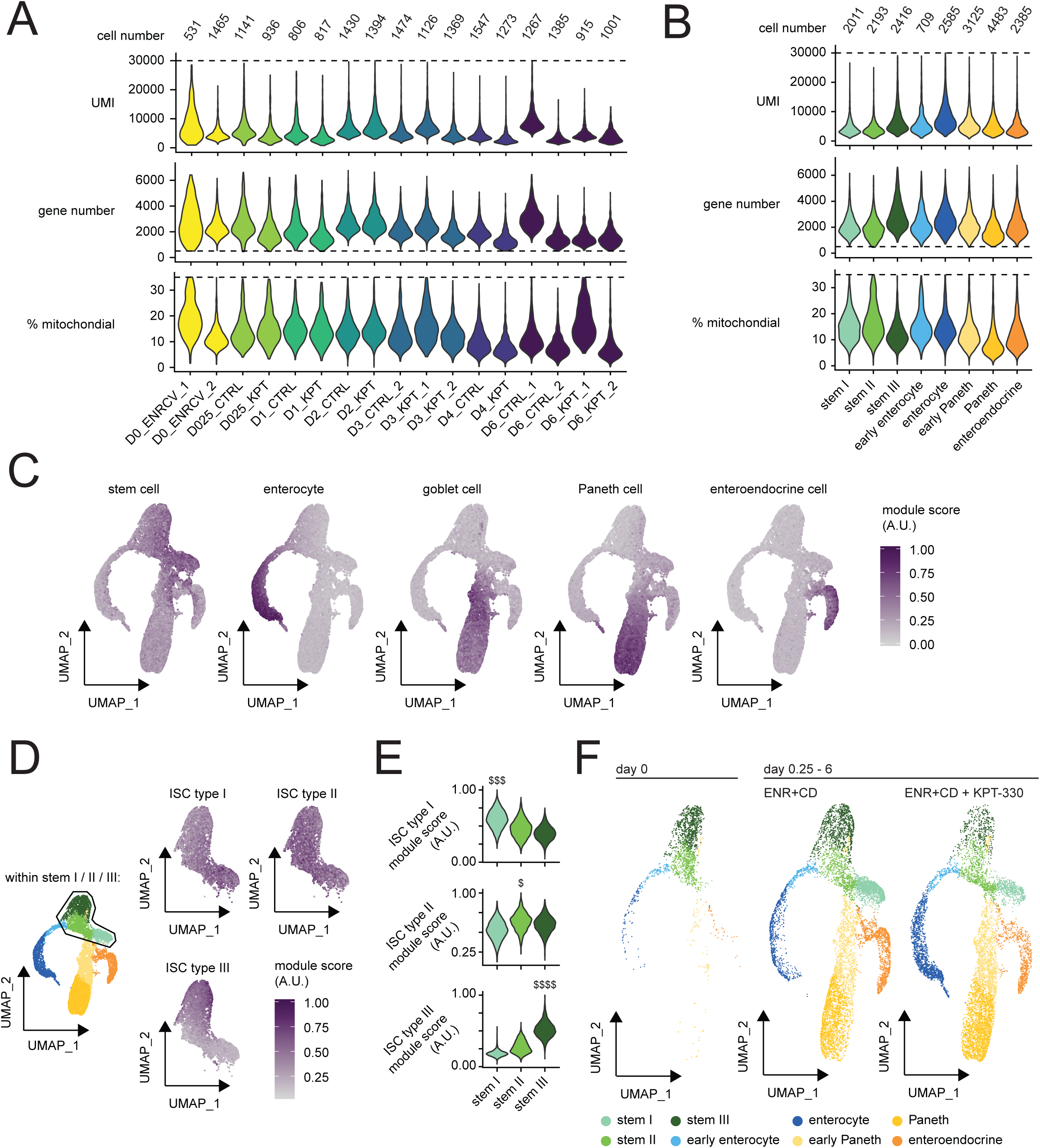
A. Single-cell RNA-seq quality metrics on a per-sample basis, including final cell number (barcodes) per array, and distributions of unique molecular identifiers per barcode (UMI), unique gene number per barcode, and percent of total UMIs corresponding to mitochondrial genes per barcode. B. Single-cell RNA-seq quality metrics on a per-cell type basis, including final cell number (barcodes) per array, and distributions of unique molecular identifiers per barcode (UMI), unique gene number per barcode, and percent of total UMIs corresponding to mitochondrial genes per barcode. C. Feature plots over organoid differentiation UMAP representing module scores derived from gene sets enriched in *in vivo* stem cells, enterocytes, goblet cells, Paneth cells, and enteroendocrine cells, each score scaled on a range from 0 to 1. D. Feature plots over organoid differentiation UMAP restricted to stem I / II / III populations representing module scores derived from gene sets enriched in *in vivo* type I / II / III intestinal stem cells (ISCs), each score scaled on a range from 0 to 1. E. Violin plots for stem I / II / III populations representing module scores derived from gene sets enriched in *in vivo* type I / II / III intestinal stem cells (ISCs), each score scaled on a range from 0 to 1. Effect size measured as Cohen’s d, $ 0.5 < d < 0.8, $$$ 1.2 < d < 2, $$$$ d > 2. F. Organoid differentiation UMAP labeled by annotated cell type, and split by day 0 (ENR+CV), day 0.25-6 control (ENR+CD), and day 0.25-6 160 nM KPT-330-treated (ENR+CD + KPT-330).

**Supplementary Figure 3.**
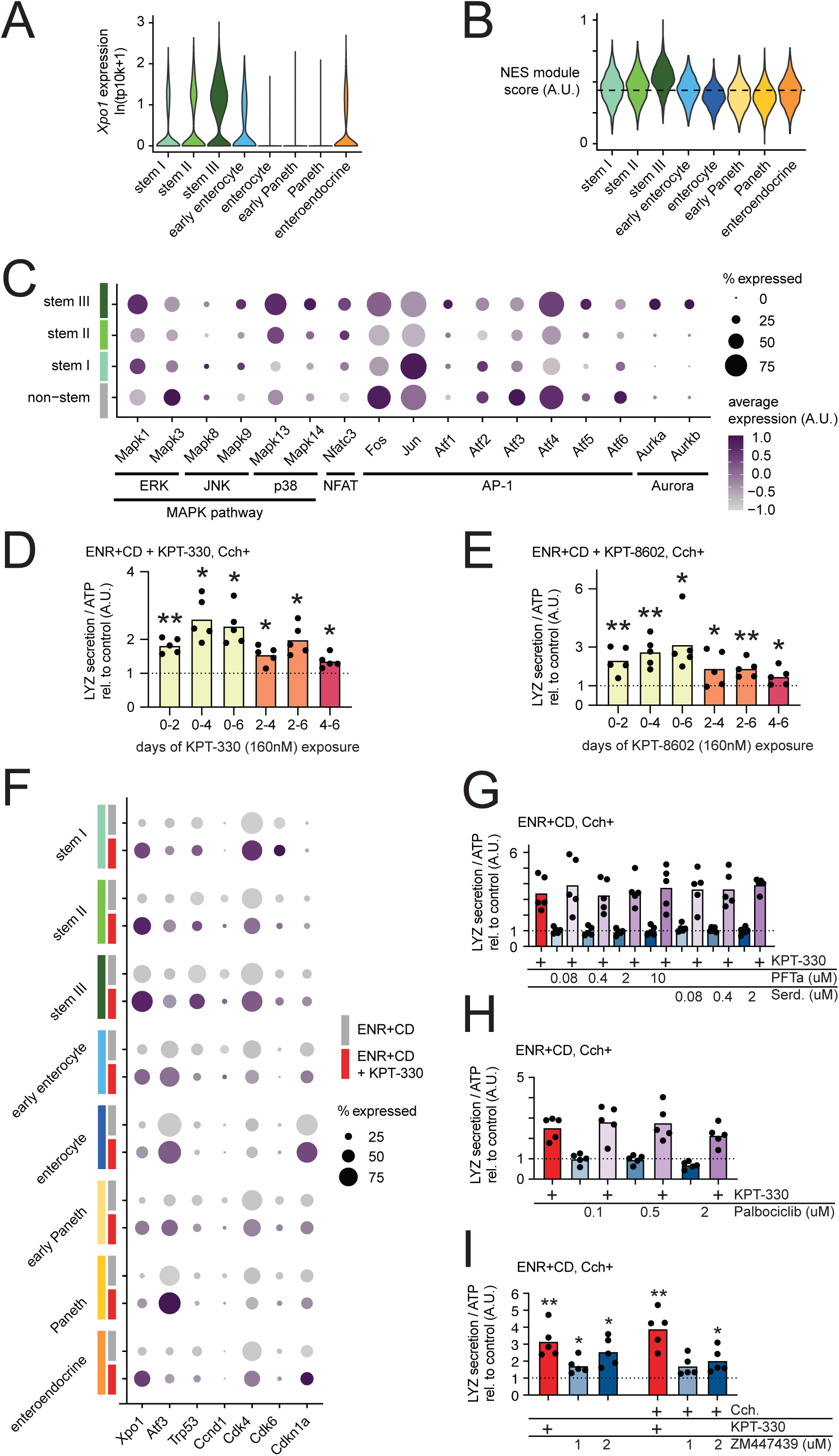
A. Single-cell RNA-seq log normalized (transcripts per 10,000 – tp10k) expression of *Xpo1* in all un-treated control cells split by cell type annotations. B. Violin plots representing module scores derived from genes with known nuclear export signals (NES) in all un-treated control cells split cell type annotations, each score scaled on a range from 0 to 1. C. Single-cell RNA-seq log normalized expression of genes involved in MAPK, NFAT, AP-1, and Aurora kinase signaling in all un-treated control cells split by non-stem, and stem I / II / III annotations. D. LYZ secretion assay for organoids treated with KPT-330 for the indicated time frame during 6 days culture in ENR+CD media. Organoids were incubated in fresh basal media with 10 μM carbachol (Cch) for 3 h on day 6. All data normalized to ATP abundance and standardized to the control in each experiment. Means and individual values are shown (N=5), dotted line represents the control value (1). One sample t-test compared to 1, followed by the two-stage linear step-up method of Benjamini, Krieger and Yekutieli for adjusting p-values; **p < 0.01, *p < 0.05. E. LYZ secretion assay for organoids treated with KPT-8602 for the indicated time frame during 6 days culture in ENR+CD media. Organoids were incubated in fresh basal media with 10 μM carbachol (Cch) for 3 h on day 6. All data normalized to ATP abundance and standardized to the control in each experiment. Means and individual values are shown (N=5), dotted line represents the control value (1). One sample t-test compared to 1, followed by the two-stage linear step-up method of Benjamini, Krieger and Yekutieli for adjusting p-values; **p < 0.01, *p < 0.05. F. Single-cell RNA-seq log normalized expression of genes known to be regulated by Xpo1 signaling between KPT-330-treated and control cells split by cell-type annotations. Color scale is relative to un-treated, purple-to-grey increasing relative-expression. G. LYZ secretion assay for organoids treated with p53 modulators, inhibitor Pifithrin-α (PFTa) and activator Serdemetan (Serd.) over 6-day culture in ENR+CD media with or without 160 nM KPT-330. Organoids were incubated in fresh basal media with 10 μM carbachol (Cch) for 3 h on day 6. All data were normalized to ATP abundance and further standardized to the control in each experiment. Means and individual values are shown (N=5), and the dotted line represents the control value (1). H. LYZ secretion assay for organoids treated with CDK4/6 inhibitor Palbociclib over 6-day culture in ENR+CD media with or without 160 nM KPT-330. Organoids were incubated in fresh basal media with 10 μM carbachol (Cch) for 3 h on day 6. All data were normalized to ATP abundance and further standardized to the control in each experiment. Means and individual values are shown (N=5), and the dotted line represents the control value (1). I. LYZ secretion assay for organoids treated with Aurora kinase inhibitor ZM447439 over 6-day culture in ENR+CD media with or without 160 nM KPT-330. Organoids were incubated in fresh basal media with 10 μM carbachol (Cch) for 3 h on day 6. All data were normalized to ATP abundance and further standardized to the control in each experiment. Means and individual values are shown (N=5), and the dotted line represents the control value (1).

**Supplementary Figure 4.**
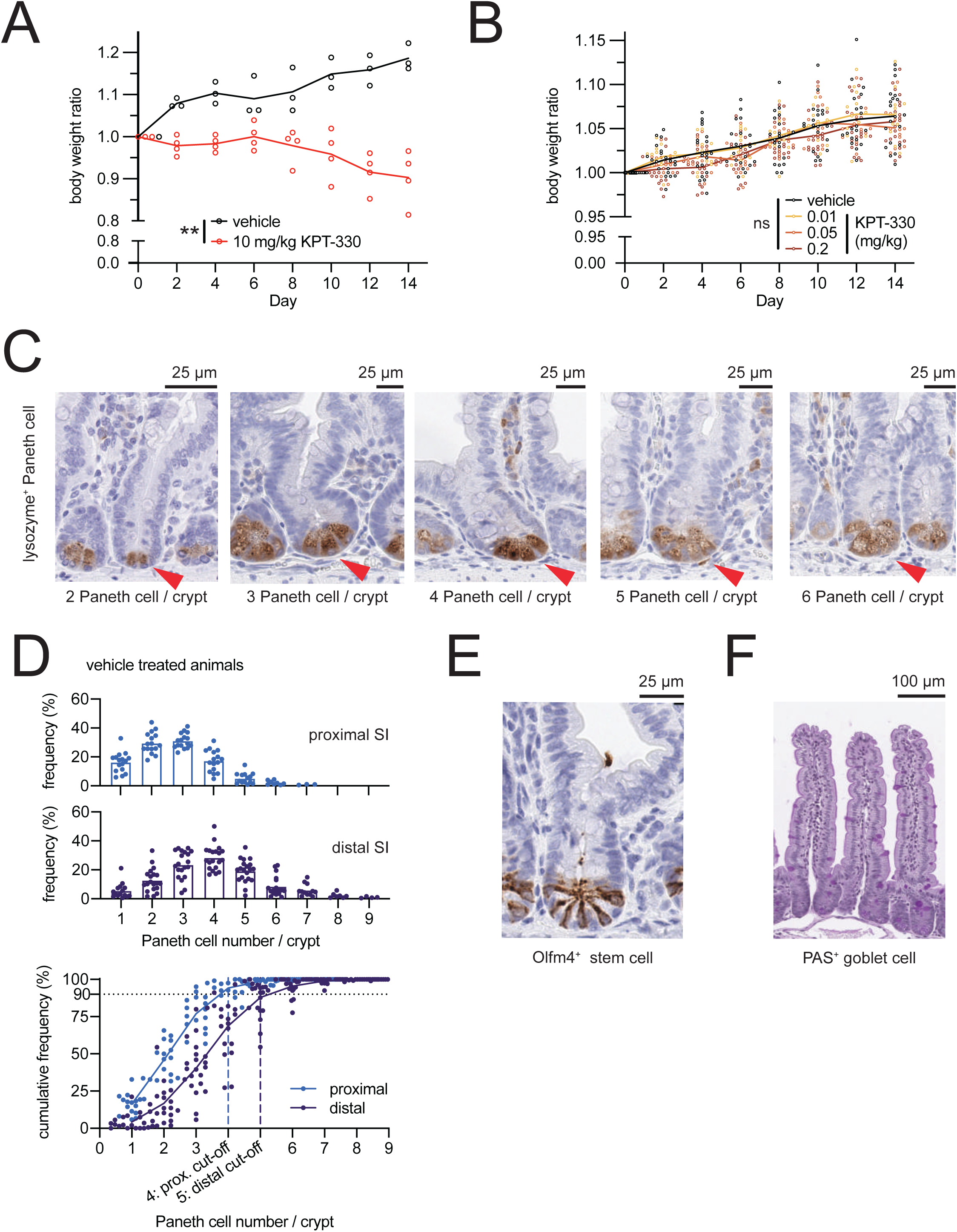
A. Animal body weight over 14-day study, normalized per-animal to day 0, of vehicle or 10 mg/kg KPT-330. Two-way ANOVA, treatment variation **p < 0.01. B. Animal body weight over 14-day study, normalized per-animal to day 0, of vehicle or 0.01, 0.05 or 0.2 mg/kg KPT-330. Two-way ANOVA, treatment variation ns p > 0.05. C. Representative histological images of small intestinal crypts with 2 to 6 Paneth cells, stained with anti-lysozyme antibody. D. Histograms of Paneth cell number in proximal or distal small intestine oh vehicle-treated animals. Cumulative frequency in proximal and distal small intestine of vehicle-treated animals was used for determining the cut-off value of Fig. 4B. E. Representative histological staining images of Olfm4^+^ stem cells in small intestine. F. Representative histological staining images of PAS^+^ goblet cells in small intestine.

## References

Arango, N.P., Yuca, E., Zhao, M., Evans, K.W., Scott, S., Kim, C., Gonzalez-Angulo, A.M., Janku, F., Ueno, N.T., Tripathy, D., Akcakanat, A., Naing, A., Meric-Bernstam, F., 2017. Selinexor (KPT-330) demonstrates anti-tumor efficacy in preclinical models of triple-negative breast cancer. Breast Cancer Res. 19, 93. doi: 10.1186/s13058-017-0878-6

Ayyaz, A., Kumar, S., Sangiorgi, B., Ghoshal, B., Gosio, J., Ouladan, S., Fink, M., Barutcu, S., Trcka, D., Shen, J., Chan, K., Wrana, J.L., Gregorieff, A., 2019. Single-cell transcriptomes of the regenerating intestine reveal a revival stem cell. Nature 569, 121–125. doi: 10.1038/s41586-019-1154-y

Azmi, A.S., Aboukameel, A., Bao, B., Sarkar, F.H., Philip, P.A., Kauffman, M., Shacham, S., Mohammad, R.M., 2013. Selective Inhibitors of Nuclear Export Block Pancreatic Cancer Cell Proliferation and Reduce Tumor Growth in Mice. Gastroenterology 144, 447–456. doi: 10.1053/j.gastro.2012.10.036

Barker, N., van Es, J.H., Kuipers, J., Kujala, P., van den Born, M., Cozijnsen, M., Haegebarth, A., Korving, J., Begthel, H., Peters, P.J., Clevers, H., 2007. Identification of stem cells in small intestine and colon by marker gene Lgr5. Nature 449, 1003–7. doi: 10.1038/nature06196

Basak, O., Beumer, J., Wiebrands, K., Seno, H., van Oudenaarden, A., Clevers, H., 2017. Induced Quiescence of Lgr5+ Stem Cells in Intestinal Organoids Enables Differentiation of Hormone-Producing Enteroendocrine Cells. Cell Stem Cell 20, 177-190.e4. doi: 10.1016/j.stem.2016.11.001

Beyaz, S., Mana, M.D., Roper, J., Kedrin, D., Saadatpour, A., Hong, S.-J., Bauer-Rowe, K.E., Xifaras, M.E., Akkad, A., Arias, E., Pinello, L., Katz, Y., Shinagare, S., Abu-Remaileh, M., Mihaylova, M.M., Lamming, D.W., Dogum, R., Guo, G., Bell, G.W., Selig, M., Nielsen, G.P., Gupta, N., Ferrone, C.R., Deshpande, V., Yuan, G.-C., Orkin, S.H., Sabatini, D.M., Yilmaz, Ö.H., 2016. High-fat diet enhances stemness and tumorigenicity of intestinal progenitors. Nature 531, 53–8. doi: 10.1038/nature17173

Biton, M., Haber, A.L., Rogel, N., Burgin, G., Beyaz, S., Schnell, A., Ashenberg, O., Su, C.-W., Smillie, C., Shekhar, K., Chen, Z., Wu, C., Ordovas-Montanes, J., Alvarez, D., Herbst, R.H., Zhang, M., Tirosh, I., Dionne, D., Nguyen, L.T., Xifaras, M.E., Shalek, A.K., von Andrian, U.H., Graham, D.B., Rozenblatt-Rosen, O., Shi, H.N., Kuchroo, V., Yilmaz, O.H., Regev, A., Xavier, R.J., 2018. T Helper Cell Cytokines Modulate Intestinal Stem Cell Renewal and Differentiation. Cell 1–14. doi: 10.1016/j.cell.2018.10.008

Cheng, C.-W., Biton, M., Haber, A.L., Gunduz, N., Eng, G., Gaynor, L.T., Tripathi, S., Calibasi-Kocal, G., Rickelt, S., Butty, V.L., Moreno-Serrano, M., Iqbal, A.M., Bauer-Rowe, K.E., Imada, S., Ulutas, M.S., Mylonas, C., Whary, M.T., Levine, S.S., Basbinar, Y., Hynes, R.O., Mino-Kenudson, M., Deshpande, V., Boyer, L.A., Fox, J.G., Terranova, C., Rai, K., Piwnica-Worms, H., Mihaylova, M.M., Regev, A., Yilmaz, Ö.H., 2019. Ketone Body Signaling Mediates Intestinal Stem Cell Homeostasis and Adaptation to Diet. Cell 178, 1115-1131.e15. doi: 10.1016/j.cell.2019.07.048

Czerniecki, S.M., Cruz, N.M., Harder, J.L., Menon, R., Annis, J., Otto, E.A., Gulieva, R.E., Islas, L. V., Kim, Y.K., Tran, L.M., Martins, T.J., Pippin, J.W., Fu, H., Kretzler, M., Shankland, S.J., Himmelfarb, J., Moon, R.T., Paragas, N., Freedman, B.S., 2018. High-Throughput Screening Enhances Kidney Organoid Differentiation from Human Pluripotent Stem Cells and Enables Automated Multidimensional Phenotyping. Cell Stem Cell 22, 929-940.e4. doi: 10.1016/j.stem.2018.04.022

De Jong, P.R., Taniguchi, K., Harris, A.R., Bertin, S., Takahashi, N., Duong, J., Campos, A.D., Powis, G., Corr, M., Karin, M., Raz, E., 2016. ERK5 signalling rescues intestinal epithelial turnover and tumour cell proliferation upon ERK1/2 abrogation. Nat. Commun. 7. doi: 10.1038/ncomms11551

Dekkers, J.F., Wiegerinck, C.L., de Jonge, H.R., Bronsveld, I., Janssens, H.M., de Winter-de Groot, K.M., Brandsma, A.M., de Jong, N.W.M., Bijvelds, M.J.C., Scholte, B.J., Nieuwenhuis, E.E.S., van den Brink, S., Clevers, H., van der Ent, C.K., Middendorp, S., Beekman, J.M., 2013. A functional CFTR assay using primary cystic fibrosis intestinal organoids. Nat. Med. 19, 939–945. doi: 10.1038/nm.3201

Draheim, K.M., Chen, H.B., Tao, Q., Moore, N., Roche, M., Lyle, S., 2010. ARRDC3 suppresses breast cancer progression by negatively regulating integrin B4. Oncogene 29, 5032–5047. doi: 10.1038/onc.2010.250

Eriguchi, Y., Takashima, S., Oka, H., Shimoji, S., Nakamura, K., Uryu, H., Shimoda, S., Iwasaki, H., Shimono, N., Ayabe, T., Akashi, K., Teshima, T., 2012. Graft-versus-host disease disrupts intestinal microbial ecology by inhibiting Paneth cell production of α-defensins. Blood 120, 223–31. doi: 10.1182/blood-2011-12-401166

Forbes, D.J., Travesa, A., Nord, M.S., Bernis, C., 2015. Nuclear transport factors: Global regulation of mitosis. Curr. Opin. Cell Biol. 35, 78–90. doi: 10.1016/j.ceb.2015.04.012

Fre, S., Huyghe, M., Mourikis, P., Robine, S., Louvard, D., Artavanis-Tsakonas, S., 2005. Notch signals control the fate of immature progenitor cells in the intestine. Nature 435, 964–8. doi: 10.1038/nature03589

Fu, S.C., Huang, H.C., Horton, P., Juan, H.F., 2013. ValidNESs: A database of validated leucine-rich nuclear export signals. Nucleic Acids Res. 41, 338–343. doi: 10.1093/nar/gks936

Gassler, N., 2017. Paneth cells in intestinal physiology and pathophysiology. World J. Gastrointest. Pathophysiol. 8, 150–160. doi: 10.4291/wjgp.v8.i4.150

Glal, D., Sudhakar, J.N., Lu, H.H., Liu, M.C., Chiang, H.Y., Liu, Y.C., Cheng, C.F., Shui, J.W., 2018. ATF3 sustains IL-22-induced STAT3 phosphorylation to maintain mucosal immunity through inhibiting phosphatases. Front. Immunol. 9. doi: 10.3389/fimmu.2018.02522

Graham, D.B., Xavier, R.J., 2020. Pathway paradigms revealed from the genetics of inflammatory bowel disease. Nature 578, 527–539. doi: 10.1038/s41586-020-2025-2

Haber, A.L., Biton, M., Rogel, N., Herbst, R.H., Shekhar, K., Smillie, C., Burgin, G., Delorey, T.M., Howitt, M.R., Katz, Y., Tirosh, I., Beyaz, S., Dionne, D., Zhang, M., Raychowdhury, R., Garrett, W.S., Rozenblatt-Rosen, O., Shi, H.N., Yilmaz, O., Xavier, R.J., Regev, A., 2017. A single-cell survey of the small intestinal epithelium. Nature. doi: 10.1038/nature24489

Hafemeister, C., Satija, R., 2019. Normalization and variance stabilization of single-cell RNA-seq data using regularized negative binomial regression. Genome Biol. 20, 296. doi: 10.1186/s13059-019-1874-1

Han, T., Schatoff, E.M., Murphy, C., Zafra, M.P., Wilkinson, J.E., Elemento, O., Dow, L.E., 2017. R-Spondin chromosome rearrangements drive Wnt-dependent tumour initiation and maintenance in the intestine. Nat. Commun. 8, 15945. doi: 10.1038/ncomms15945

Hayase, E., Hashimoto, D., Nakamura, K., Noizat, C., Ogasawara, R., Takahashi, S., Ohigashi, H., Yokoi, Y., Sugimoto, R., Matsuoka, S., Ara, T., Yokoyama, E., Yamakawa, T., Ebata, K., Kondo, T., Hiramine, R., Aizawa, T., Ogura, Y., Hayashi, T., Mori, H., Kurokawa, K., Tomizuka, K., Ayabe, T., Teshima, T., 2017. R-Spondin1 expands Paneth cells and prevents dysbiosis induced by graft-versus-host disease. J. Exp. Med. 214, 3507–3518. doi: 10.1084/jem.20170418

Heuberger, J., Kosel, F., Qi, J., Grossmann, K.S., Rajewsky, K., Birchmeier, W., 2014. Shp2/MAPK signaling controls goblet/paneth cell fate decisions in the intestine. Proc. Natl. Acad. Sci. U. S. A. 111, 3472–3477. doi: 10.1073/pnas.1309342111

Hing, Z.A., Fung, H.Y.J., Ranganathan, P., Mitchell, S., El-Gamal, D., Woyach, J.A., Williams, K., Goettl, V.M., Smith, J., Yu, X., Meng, X., Sun, Q., Cagatay, T., Lehman, A.M., Lucas, D.M., Baloglu, E., Shacham, S., Kauffman, M.G., Byrd, J.C., Chook, Y.M., Garzon, R., Lapalombella, R., 2016. Next-generation XPO1 inhibitor shows improved efficacy and in vivo tolerability in hematological malignancies. Leukemia 30, 2364–2372. doi: 10.1038/leu.2016.136

Hughes, T.K., Wadsworth, M.H., Gierahn, T.M., Do, T., Weiss, D., Andrade, P.R., Ma, F., Silva, B.J. de A., Shao, S., Tsoi, L.C., Ordovas-Montanes, J., Gudjonsson, J.E., Modlin, R.L., Love, J.C., Shalek, A.K., 2019. Highly Efficient, Massively-Parallel Single-Cell RNA-Seq Reveals Cellular States and Molecular Features of Human Skin Pathology. bioRxiv 689273. doi: 10.1101/689273

Ireland, H., Houghton, C., Howard, L., Winton, D.J., 2005. Cellular inheritance of a Cre-activated reporter gene to determine paneth cell longevity in the murine small intestine. Dev. Dyn. 233, 1332–1336. doi: 10.1002/dvdy.20446

Jadhav, K., Zhang, Y., 2017. Activating transcription factor 3 in immune response and metabolic regulation. Liver Res. 1, 96–102. doi: 10.1016/j.livres.2017.08.001

Jensen, J., Pedersen, E.E., Galante, P., Hald, J., Heller, R.S., Ishibashi, M., Kageyama, R., Guillemot, F., Serup, P., Madsen, O.D., 2000. Control of endodermal endocrine development by Hes-1. Nat. Genet. 24, 36–44. doi: 10.1038/71657

Khor, B., Gardet, A., Xavier, R.J., 2011. Genetics and pathogenesis of inflammatory bowel disease. Nature 474, 307–17. doi: 10.1038/nature10209

Kim, K.-A., Kakitani, M., Zhao, J., Oshima, T., Tang, T., Binnerts, M., Liu, Y., Boyle, B., Park, E., Emtage, P., Funk, W.D., Tomizuka, K., 2005. Mitogenic influence of human R-spondin1 on the intestinal epithelium. Science 309, 1256–9. doi: 10.1126/science.1112521

Korving, J., Moll, J., Voest, E.E., Weeber, F., de Ligt, J., Rottenberg, S., Bounova, G., Boj, S.F., Kopper, O., Vries, R.G.J., van de Ven, M., Wessels, L., Nijman, I.J., Lelieveld, D., Ernst, R.F., Gogola, E., Blokzijl, F., Sachs, N., Duarte, A.A., Cuppen, E., Gracanin, A., Hoogstraat, M., van Hoeck, A., Clevers, H., Zinzalla, V., van Boxtel, R., Begthel, H., Egan, D.A., Balgobind, A.V., Wind, K., van Diest, P.J., 2017. A Living Biobank of Breast Cancer Organoids Captures Disease Heterogeneity. Cell 172, 373-386.e10. doi: 10.1016/j.cell.2017.11.010

Langhans, S.A., 2018. Three-Dimensional in Vitro Cell Culture Models in Drug Discovery and Drug Repositioning. Front. Pharmacol. 9, 1–14. doi: 10.3389/fphar.2018.00006

Liberzon, A., Birger, C., Thorvaldsdóttir, H., Ghandi, M., Mesirov, J.P., Tamayo, P., 2015. The Molecular Signatures Database Hallmark Gene Set Collection. Cell Syst. 1, 417–425. doi: 10.1016/j.cels.2015.12.004

Liu, T.-C., Gurram, B., Baldridge, M.T., Head, R., Lam, V., Luo, C., Cao, Y., Simpson, P., Hayward, M., Holtz, M.L., Bousounis, P., Noe, J., Lerner, D., Cabrera, J., Biank, V., Stephens, M., Huttenhower, C., McGovern, D.P.B., Xavier, R.J., Stappenbeck, T.S., Salzman, N.H., 2016. Paneth cell defects in Crohn’s disease patients promote dysbiosis. JCI insight 1, e86907. doi: 10.1172/jci.insight.86907

Macosko, E.Z., Basu, A., Satija, R., Nemesh, J., Shekhar, K., Goldman, M., Tirosh, I., Bialas, A.R., Kamitaki, N., Martersteck, E.M., Trombetta, J.J., Weitz, D.A., Sanes, J.R., Shalek, A.K., Regev, A., McCarroll, S.A., 2015. Highly parallel genome-wide expression profiling of individual cells using nanoliter droplets. Cell 161, 1202–1214. doi: 10.1016/j.cell.2015.05.002

McElroy, S.J., Underwood, M.A., Sherman, M.P., 2013. Paneth cells and necrotizing enterocolitis: a novel hypothesis for disease pathogenesis. Neonatology 103, 10–20. doi: 10.1159/000342340

McGinnis, C.S., Murrow, L.M., Gartner, Z.J., 2019. DoubletFinder: Doublet Detection in Single-Cell RNA Sequencing Data Using Artificial Nearest Neighbors. Cell Syst. 8, 329-337.e4. doi: 10.1016/j.cels.2019.03.003

McGuckin, M. a, Eri, R., Simms, L. a, Florin, T.H.J., Radford-Smith, G., 2009. Intestinal barrier dysfunction in inflammatory bowel diseases. Inflamm. Bowel Dis. 15, 100–13. doi: 10.1002/ibd.20539

Mead, B.E., Karp, J.M., 2019. All models are wrong, but some organoids may be useful. Genome Biol. 20, 66. doi: 10.1186/s13059-019-1677-4

Mead, B.E., Ordovas-Montanes, J., Braun, A.P., Levy, L.E., Bhargava, P., Szucs, M.J., Ammendolia, D.A., MacMullan, M.A., Yin, X., Hughes, T.K., Wadsworth, M.H., Ahmad, R., Rakoff-Nahoum, S., Carr, S.A., Langer, R., Collins, J.J., Shalek, A.K., Karp, J.M., 2018. Harnessing single-cell genomics to improve the physiological fidelity of organoid-derived cell types. BMC Biol. 16, 62. doi: 10.1186/s12915-018-0527-2

Mpindi, J.P., Swapnil, P., Dmitrii, B., Jani, S., Saeed, K., Wennerberg, K., Aittokallio, T., Östling, P., Kallioniemi, O., 2015. Impact of normalization methods on high-throughput screening data with high hit rates and drug testing with dose-response data. Bioinformatics 31, 3815–3821. doi: 10.1093/bioinformatics/btv455

Naik, S., Larsen, S.B., Gomez, N.C., Alaverdyan, K., Sendoel, A., Yuan, S., Polak, L., Kulukian, A., Chai, S., Fuchs, E., 2017. Inflammatory memory sensitizes skin epithelial stem cells to tissue damage. Nature 550, 475–480. doi: 10.1038/nature24271

Okubo, T., Hogan, B.L.M., 2004. Hyperactive Wnt signaling changes the developmental potential of embryonic lung endoderm. J. Biol. 3, 11. doi: 10.1186/jbiol3

Ordovas-Montanes, J., Dwyer, D.F., Nyquist, S.K., Buchheit, K.M., Vukovic, M., Deb, C., Wadsworth, M.H., Hughes, T.K., Kazer, S.W., Yoshimoto, E., Cahill, K.N., Bhattacharyya, N., Katz, H.R., Berger, B., Laidlaw, T.M., Boyce, J.A., Barrett, N.A., Shalek, A.K., 2018. Allergic inflammatory memory in human respiratory epithelial progenitor cells. Nature 560, 649–654. doi: 10.1038/s41586-018-0449-8

Pinto, D., Gregorieff, A., Begthel, H., Clevers, H., 2003. Canonical Wnt signals are essential for homeostasis of the intestinal epithelium. Genes Dev. 17, 1709–1713. doi: 10.1101/gad.267103

Roulis, M., Kaklamanos, A., Schernthanner, M., Bielecki, P., Zhao, J., Kaffe, E., Frommelt, L.-S., Qu, R., Knapp, M.S., Henriques, A., Chalkidi, N., Koliaraki, V., Jiao, J., Brewer, J.R., Bacher, M., Blackburn, H.N., Zhao, X., Breyer, R.M., Aidinis, V., Jain, D., Su, B., Herschman, H.R., Kluger, Y., Kollias, G., Flavell, R.A., 2020. Paracrine orchestration of intestinal tumorigenesis by a mesenchymal niche. Nature. doi: 10.1038/s41586-020-2166-3

Sansom, O.J., Reed, K.R., Hayes, A.J., Ireland, H., Brinkmann, H., Newton, I.P., Batlle, E., Simon-Assmann, P., Clevers, H., Nathke, I.S., Clarke, A.R., Winton, D.J., 2004. Loss of Apc in vivo immediately perturbs Wnt signaling, differentiation, and migration. Genes Dev. 18, 1385–90. doi: 10.1101/gad.287404

Sato, T., Vries, R.G., Snippert, H.J., van de Wetering, M., Barker, N., Stange, D.E., van Es, J.H., Abo, A., Kujala, P., Peters, P.J., Clevers, H., 2009. Single Lgr5 stem cells build crypt-villus structures in vitro without a mesenchymal niche. Nature 459, 262–5. doi: 10.1038/nature07935

Sendino, M., Omaetxebarria, M.J., Rodríguez, J.A., 2018. Hitting a moving target: inhibition of the nuclear export receptor XPO1/CRM1 as a therapeutic approach in cancer. Cancer Drug Resist. doi: 10.20517/cdr.2018.09

Serra, D., Mayr, U., Boni, A., Lukonin, I., Rempfler, M., Challet Meylan, L., Stadler, M.B., Strnad, P., Papasaikas, P., Vischi, D., Waldt, A., Roma, G., Liberali, P., 2019. Self-organization and symmetry breaking in intestinal organoid development. Nature. doi: 10.1038/s41586-019-1146-y

Sherman, M.P., Bennett, S.H., Hwang, F.F.Y., Sherman, J., Bevins, C.L., 2005. Paneth cells and antibacterial host defense in neonatal small intestine. Infect. Immun. 73, 6143–6. doi: 10.1128/IAI.73.9.6143-6146.2005

Subramanian, A., Tamayo, P., Mootha, V.K., Mukherjee, S., Ebert, B.L., Gillette, M.A., Paulovich, A., Pomeroy, S.L., Golub, T.R., Lander, E.S., Mesirov, J.P., 2005. Gene set enrichment analysis: A knowledge-based approach for interpreting genome-wide expression profiles. Proc. Natl. Acad. Sci. 102, 15545–15550. doi: 10.1073/pnas.0506580102

Sun, Q., Chen, X., Zhou, Q., Burstein, E., Yang, S., Jia, D., 2016. Inhibiting cancer cell hallmark features through nuclear export inhibition. Signal Transduct. Target. Ther. 1, 34–36. doi: 10.1038/sigtrans.2016.10

Tanner, S.M., Berryhill, T.F., Ellenburg, J.L., Jilling, T., Cleveland, D.S., Lorenz, R.G., Martin, C.A., 2015. Pathogenesis of necrotizing enterocolitis: modeling the innate immune response. Am. J. Pathol. 185, 4–16. doi: 10.1016/j.ajpath.2014.08.028

Tetteh, P.W., Basak, O., Farin, H.F., Wiebrands, K., Kretzschmar, K., Begthel, H., van den Born, M., Korving, J., de Sauvage, F., van Es, J.H., van Oudenaarden, A., Clevers, H., 2016. Replacement of Lost Lgr5-Positive Stem Cells through Plasticity of Their Enterocyte-Lineage Daughters. Cell Stem Cell 18, 203–213. doi: 10.1016/j.stem.2016.01.001

Tirosh, I., Izar, B., Prakadan, S.M., Wadsworth, M.H., Treacy, D., Trombetta, J.J., Rotem, A., Rodman, C., Lian, C., Murphy, G., Fallahi-Sichani, M., Dutton-Regester, K., Lin, J.-R.R., Cohen, O., Shah, P., Lu, D., Genshaft, A.S., Hughes, T.K., Ziegler, C.G.K.K., Kazer, S.W., Gaillard, A., Kolb, K.E., Villani, A.-C.C., Johannessen, C.M., Andreev, A.Y., Van Allen, E.M., Bertagnolli, M., Sorger, P.K., Sullivan, R.J., Flaherty, K.T., Frederick, D.T., Jané-Valbuena, J., Yoon, C.H., Rozenblatt-Rosen, O., Shalek, A.K., Regev, A., Garraway, L.A., 2016. Dissecting the multicellular ecosystem of metastatic melanoma by single-cell RNA-seq. Science 352, 189–96. doi: 10.1126/science.aad0501

Tyler, P.M., Servos, M.M., de Vries, R.C., Klebanov, B., Kashyap, T., Sacham, S., Landesman, Y., Dougan, M., Dougan, S.K., 2017. Clinical Dosing Regimen of Selinexor Maintains Normal Immune Homeostasis and T-cell Effector Function in Mice: Implications for Combination with Immunotherapy. Mol. Cancer Ther. 16, 428–439. doi: 10.1158/1535-7163.MCT-16-0496

VanDussen, K.L., Carulli, A.J., Keeley, T.M., Patel, S.R., Puthoff, B.J., Magness, S.T., Tran, I.T., Maillard, I., Siebel, C., Kolterud, A., Grosse, A.S., Gumucio, D.L., Ernst, S.A., Tsai, Y.-H., Dempsey, P.J., Samuelson, L.C., 2012. Notch signaling modulates proliferation and differentiation of intestinal crypt base columnar stem cells. Development 139, 488–497. doi: 10.1242/dev.070763

VanDussen, K.L., Samuelson, L.C., 2010. Mouse atonal homolog 1 directs intestinal progenitors to secretory cell rather than absorptive cell fate. Dev. Biol. 346, 215–23. doi: 10.1016/j.ydbio.2010.07.026

Väremo, L., Nielsen, J., Nookaew, I., 2013. Enriching the gene set analysis of genome-wide data by incorporating directionality of gene expression and combining statistical hypotheses and methods. Nucleic Acids Res. 41, 4378–4391. doi: 10.1093/nar/gkt111

von Moltke, J., Ji, M., Liang, H., Locksley, R.M., 2016. Tuft-cell-derived IL-25 regulates an intestinal ILC2–epithelial response circuit. Nature 529, 221–225. doi: 10.1038/nature16161

Wang, A.Y., Liu, H., 2019. The past, present, and future of CRM1/XPO1 inhibitors. Stem Cell Investig. 6. doi: 10.21037/sci.2019.02.03

White, J.R., Gong, H., Pope, B., Schlievert, P., McElroy, S.J., 2017. Paneth-cell-disruption-induced necrotizing enterocolitis in mice requires live bacteria and occurs independently of TLR4 signaling. Dis. Model. Mech. 10, 727–736. doi: 10.1242/dmm.028589

Wu, A., Yu, B., Zhang, K., Xu, Z., Wu, D., He, J., Luo, J., Luo, Y., Yu, J., Zheng, P., Che, L., Mao, X., Huang, Z., Wang, L., Zhao, J., Chen, D., 2020. Transmissible gastroenteritis virus targets Paneth cells to inhibit the self-renewal and differentiation of Lgr5 intestinal stem cells via Notch signaling. Cell Death Dis. 11, 40. doi: 10.1038/s41419-020-2233-6

Xavier, R.J., Podolsky, D.K., 2007. Unravelling the pathogenesis of inflammatory bowel disease. Nature 448, 427–34. doi: 10.1038/nature06005

Yan, K.S., Gevaert, O., Zheng, G.X.Y., Anchang, B., Probert, C.S., Larkin, K.A., Davies, P.S., Cheng, Z., Kaddis, J.S., Han, A., Roelf, K., Calderon, R.I., Cynn, E., Hu, X., Mandleywala, K., Wilhelmy, J., Grimes, S.M., Corney, D.C., Boutet, S.C., Terry, J.M., Belgrader, P., Ziraldo, S.B., Mikkelsen, T.S., Wang, F., von Furstenberg, R.J., Smith, N.R., Chandrakesan, P., May, R., Chrissy, M.A.S., Jain, R., Cartwright, C.A., Niland, J.C., Hong, Y.-K., Carrington, J., Breault, D.T., Epstein, J., Houchen, C.W., Lynch, J.P., Martin, M.G., Plevritis, S.K., Curtis, C., Ji, H.P., Li, L., Henning, S.J., Wong, M.H., Kuo, C.J., 2017. Intestinal Enteroendocrine Lineage Cells Possess Homeostatic and Injury-Inducible Stem Cell Activity. Cell Stem Cell 21, 78-90.e6. doi: 10.1016/j.stem.2017.06.014

Yang, H.W., Chung, M., Kudo, T., Meyer, T., 2017. Competing memories of mitogen and p53 signalling control cell-cycle entry. Nature 549, 404–408. doi: 10.1038/nature23880

Yin, X., Farin, H.F., van Es, J.H., Clevers, H., Langer, R., Karp, J.M., 2014. Niche-independent high-purity cultures of Lgr5+ intestinal stem cells and their progeny. Nat. Methods 11, 106–12. doi: 10.1038/nmeth.2737

Yousefi, M., Li, L., Lengner, C.J., 2017. Hierarchy and Plasticity in the Intestinal Stem Cell Compartment. Trends Cell Biol. 27, 753–764. doi: 10.1016/j.tcb.2017.06.006

Zhang, X.D., 2011. Hit Selection in Genome-Scale RNAi Screens with Replicates, in: Optimal High-Throughput Screening. Cambridge University Press, Cambridge, pp. 83–108. doi: 10.1017/CBO9780511973888.007

Zheng, Y., Gery, S., Sun, H., Shacham, S., Kauffman, M., Koeffler, H.P., 2014. KPT-330 inhibitor of XPO1-mediated nuclear export has anti-proliferative activity in hepatocellular carcinoma. Cancer Chemother. Pharmacol. 74, 487–495. doi: 10.1007/s00280-014-2495-8

Zhou, J., Edgar, B.A., Boutros, M., 2017. ATF3 acts as a rheostat to control JNK signalling during intestinal regeneration. Nat. Commun. 8, 1–15. doi: 10.1038/ncomms14289

